# ProtWave-VAE: Integrating autoregressive sampling with latent-based inference for data-driven protein design

**DOI:** 10.1101/2023.04.23.537971

**Authors:** Niksa Praljak, Xinran Lian, Rama Ranganathan, Andrew L. Ferguson

## Abstract

Deep generative models (DGMs) have shown great success in the understanding of data-driven design of proteins. Variational autoencoders (VAEs) are a popular DGM approach that can learn the correlated patterns of amino acid mutations within a multiple sequence alignment (MSA) of protein sequences and distill this information into a low-dimensional latent space to expose phylogenetic and functional relationships and guide generative protein design. Autoregressive (AR) models are another popular DGM approach that typically lack a low-dimensional latent embedding but do not require training sequences to be aligned into an MSA and enable the design of variable length proteins. In this work, we propose ProtWave-VAE as a novel and lightweight DGM employing an information maximizing VAE with a dilated convolution encoder and autoregressive WaveNet decoder. This architecture blends the strengths of the VAE and AR paradigms in enabling training over unaligned sequence data and the conditional generative design of variable length sequences from an interpretable low-dimensional learned latent space. We evaluate the model’s ability to infer patterns and design rules within alignment-free homologous protein family sequences and to design novel synthetic proteins in four diverse protein families. We show that our model can infer meaningful functional and phylogenetic embeddings within latent spaces and make highly accurate predictions within semi-supervised downstream fitness prediction tasks. In an application to the C-terminal SH3 domain in the Sho1 transmembrane osmosensing receptor in baker’s yeast, we subject ProtWave-VAE designed sequences to experimental gene synthesis and select-seq assays for osmosensing function to show that the model enables *de novo* generative design, conditional C-terminus diversification, and engineering of osmosensing function into SH3 paralogs.

## Introduction

A long-standing goal in protein engineering and chemistry has been the design of novel synthetic proteins with engineered function and properties. Natural proteins have evolved under genetic drift and natural selection to robustly perform complex functions, such as ligand binding, molecular recognition, and substrate specific catalysis. With the exponential growth of sequenced protein datasets and the advent of mature deep learning models, modern machine learning tools have become ubiquitous in the engineering of novel proteins with desired functions.^1^ In particular, deep generative models (DGMs) offer a powerful modeling paradigm to learn sequence-to-function mappings and employ these relations for synthetic protein design.^2–5^ Two primary DGM paradigms have demonstrated substantial success in protein engineering: autoregressive (AR) language models^6–11^ and variational autoencoders (VAEs).^12–15,15–20^ AR models can operate on variable length sequences meaning that they do not require the construction of multiple sequence alignments and can be used to learn and generate novel sequences with high variability and diverse lengths.^7,8^ Since protein sequences from non-homologous families or within homologous families with high variability present challenges in constructing alignments,^7^ AR generative models are well-suited for alignment-free training, prediction, and design. AR models have been shown to successfully engineer novel nanobodies by designing the complementary determining region 3,^7^ demonstrate competitive performance in mutation and contact prediction, ^21^ and have been used to design functional synthetic proteins across diverse protein families.^8^ A limitation of autoregressive models is their typical lack of a low-dimensional learned latent space that exposes interpretable phylogenetic and functional relationships and can be used to guide conditional generation of synthetic sequences.

In contrast, VAEs naturally infer a latent space that is subsequently used for conditional generation of novel sequences. VAE models have been shown to accurately predict single-mutant effects,^13^ infer a homologous family’s phylogeny and fitness within the latent space,^12^ design *in vivo* signaling ortholog proteins, ^14^ and diversify synthetic AAV capsids.^22^ Although these models possess the attractive capability to infer a biologically meaningful latent space, VAEs are prone to posterior collapse (i.e., the decoder ignores latent space and the latent space fails to learn a useful representation of the training data), which can limit the use of powerful and expressive decoders for generative sequence design.^23–27^ This means that VAEs can struggle to incorporate autoregressive decoders for generating variable-length synthetic sequences and training over alignment-free homologous protein datasets.

In this work, we propose ProtWave-VAE as a new DGM architecture integrating the desirable features of the AR and VAE paradigms to permit learning of an interpretable low-dimensional latent space for conditional sequence generation and the ability to learn over non-homologous, unaligned sequence data and generate variable-length protein sequences (Figure 1). To overcome posterior collapse^23^ and enable the use of expressive AR VAE decoders, we incorporated an Information Maximizing (InfoMax) loss objective instead of the common ELBO training objective.^24^ The InfoMax loss is similar to ELBO, but, pre-factor weights and additional max-mean discrepancy regularization terms are introduced to motivate better inference and regularization. A mutual information maximization term is introduced to encourage high mutual information between the input vectors and latent space embeddings. We implement a WaveNet-based autoregressive generator^28^ for our decoder that avoids vanishing or exploding gradients by using dilated causal convolutions. Previously, models have been developed that combine VAEs with dilated causal convolutions as the decoder for text generation,^27^ but this approach has yet to be explored for protein design and fitness prediction. These convolutions are much faster than recurrent networks during training time, offer superior inference of long-range correlations, and are computationally less expensive than large-scale language models. We note that the AR-VAE model can be modularly extended to use other powerful and expressive decoders, such as autoregressive transformer-based decoders.^29^

**Figure 1:**
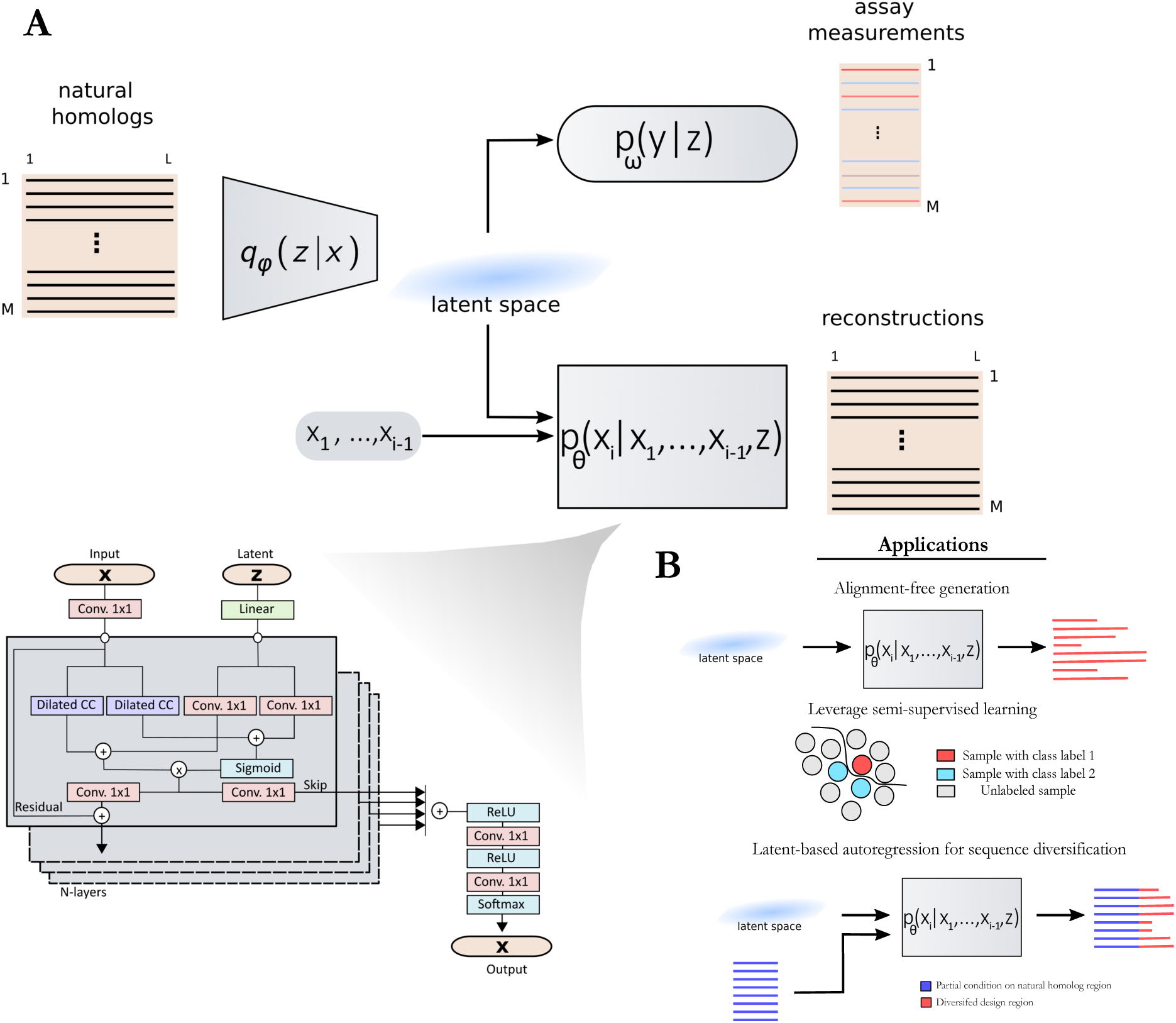
(A) Schematic illustration of the ProtWave-VAE model integrating an InfoMax VAE with convolutional encoder and WaveNet autoregressive decoder. This unsupervised learning model may be made semi-supervised by incorporating an optional top model comprising, a discriminative multi-layer perception to regress function on the protein location within the latent space embedding. The model architecture employs a gated dilated convolutional encoder *q_ϕ_*(*z|x*), a WaveNet (i.e., gated dilated causal convolution) autoregressive decoder *p_θ_*(*x_i_|x*_1_*, …, x_i−_*_1_*, z*), and a supervised neural regression model *p_ω_*(*y|z*) to predict functional assay measurements when available. (B) By combining latent inference and autoregressive generation, our model enables (i) alignment-free inference and variable-length generation, (ii) semi-supervised learning, and (iii) conditional generation based on N-terminal residues and latent space conditioning vectors. The first application allows training models on protein families that require no multiple-sequence alignments (MSAs). The second application provides and leverages assay measurements during the generative learning process and reshapes the latent space for better control of the functional design of proteins. The third application permits sequence diversification by conditioning on an N-terminus sequence motif of a natural homolog and conditioning on latent embeddings to generate and inpaint the C-terminus region. This final application is not restricted to the C-terminus diversification of natural homologs; rather, it can also be utilized by conditioning protein tags (such as expression-tags or affinity-tags) and filling in missing protein sequences through inpainting.

The ProtWave-VAE shares similarities with, but is differentiated from, a number of related approaches in the literature. The ProT-VAE model of Sevgen et al.^19^ uses a VAE architecture employing a large-scale pre-trained ProtT5 encoder and decoder and has shown substantial promise for alignment-free protein design. ProtWave-VAE is distinguished by its incorporation of latent conditioning and autoregressive sampling along the decoder path, and its lightweight architecture comprising *∼*10^6^-10^7^ trainable parameters relative to *∼*10^9^ for ProT-VAE. Prior work using WaveNet autoregressive models demonstrated competitive prediction of mutant effects and the design novel nanobodies,^7^ but the absence of a latent space precluded these approaches from leveraging latent conditioning for controlled protein generation. By virtue of the variational inference of meaningful biological latent codes, latent conditioning is available to ProtWave-VAE. Similar to other autoregressive models for protein design which leverage a large transformer (ProGen1^8^) and LSTM architecture (UniRep^6^), our approach infers a regularized latent space for meaningful conditional information for the autoregressive decoder. Hawkins et al.^18^ have previously demonstrated the use of an AR-VAE model for protein design, but our model employs a powerful WaveNet decoder instead of GRU enabled by our use of an Information Maximizing VAE loss to prevent posterior collapse.^24^

We demonstrate and test the ProtWave-VAE in applications to both retrospective protein function prediction tasks and *de novo* protein design wherein the sequences designed by the model are experimentally synthesized and tested. First, we show that our approach can infer biologically meaningful latent spaces while incorporating an expressive autoregressive generator and learning on alignment-free sequences for four selected protein families.^30^ To assess the generative capacity of our model to synthesize well-folded tertiary structures, we sampled novel sequences and predicted their tertiary structures using AlphaFold2.^31,32^ Despite not being exposed to any structural information during training, we find that the predicted structures of the synthetic sequences recapitulate the native folds of the corresponding to the protein family. Second, we extend the training objective to a semi-supervised learning paradigm^33^ for fitness landscape prediction and benchmark our semi-supervised model variant on four fitness landscape predictions within TAPE and FLIP protein function prediction tasks.^34,35^ Our model predictions are competitive or superior to with current state-of-the-art approaches employing models typically using approximately an order of magnitude more trainable parameters. Third, using the AroQ Chorismate mutase family with fitness assay measurements^36^ as a training set, we demonstrate that our model can reshape the latent space to induce functional gradients valuable for conditional generation of novel synthetic proteins with elevated functionality without losing generative capabilities. Fourth, we introduce a method to perform C-terminal diversification of natural protein sequences by conditioning on a user-specified number of N-terminal residues and a latent space conditioning vectors. This enables us to introduce sequence diversity into the C-terminal region of the sequence while also engineering in desired phylogeny and/or function through latent space conditioning. We demonstrate this approach in applications to homologs of the CM protein. Fifth, to experimentally verify our model’s generative capacity, we used a high-throughput *in vivo* select-seq to measure the binding of the Src homology 3 (SH3) domain in *Saccharomyces cerevisiae* (i.e. Baker’s yeast) to its cognate pbs2 ligand within an osmosensing pathway that protects the cell from high salt conditions by activating a homeostatic response. ^14,37^ Using this assay, we generated novel sequences that rescue osmosensing function and partially diversified natural SH3 homologs that maintain or elevate functionality. In summary, the ProtWave-VAE model presents a novel AR-VAE DGM to learn informative and meaningful low-dimensional latent space embeddings from unaligned training sequences, and permit the conditional autoregressive generation of variable-length synthetic sequence with engineered N-terminal residues and/or latent vectors informing desired phylogeny and/or function.

## Results and Discussion

### Alignment-free learning of latent space embeddings

Our first test of the model was to assess the degree to which the latent space can expose biologically meaningful representations of phylogenetic and functional patterns in unaligned homologous protein data sets. Identifying these “design rules” (i.e., correlated patterns of amino acid mutations underpinning phylogeny and function) is a pre-requisite to tailored design of synthetic functional proteins.^15^ To do so, we trained independent ProtWave-VAE models over four protein families:^30^ G-protein (GTPase, Ras-like; PFAM PF00071), Dihydrofolate reductase (DHFR; PFAM PF00186), beta-lactamase (PFAM PF13354), and S1A serine protease.^38^ Full details of model training and hyperparameter optimization, including latent space dimensionality, are provided in the Materials and Methods. For visual clarity and consistency, irrespective of the latent space dimensionality, we present 2D latent space projections into the top two principal components identified by principal components analysis (PCA). By annotating the PCA projections of the latent embeddings with known phylogenetic and functional information for the natural homologs, we find that in all four cases the ProtWave-VAE learns to disentangle the training sequences into their phylogenetic groups and functional sub-classes (Figure 2A and S2A). For G-protein, the model inferred disentangled representations in terms of functional sub-classes – Ras, Rab, Rac, Rho – and phylogeny – Metazoa, Amoebozoa, Viridiplantae, Fungi, Alveolata (Figure 2A-i). For DHFR, the latent embeddings show well-defined clusters annotated by phylogenic groups – Eukaryota, Firmicutes, Actinobacteria (Figure 2A-ii). Similarly, for the lactamase family, the PCA projections of the latent space show disentangled phylogenic groups (Figure S2A-ii). Interestingly, for the S1A family, the model can infer meaningful representations in terms of functional specificity – Trypsin versus Chymotrypsin – and homologs by their corresponding species information – vertebrate or invertebrate species and warm or cold environment species (Figure S2A-i).

**Figure 2:**
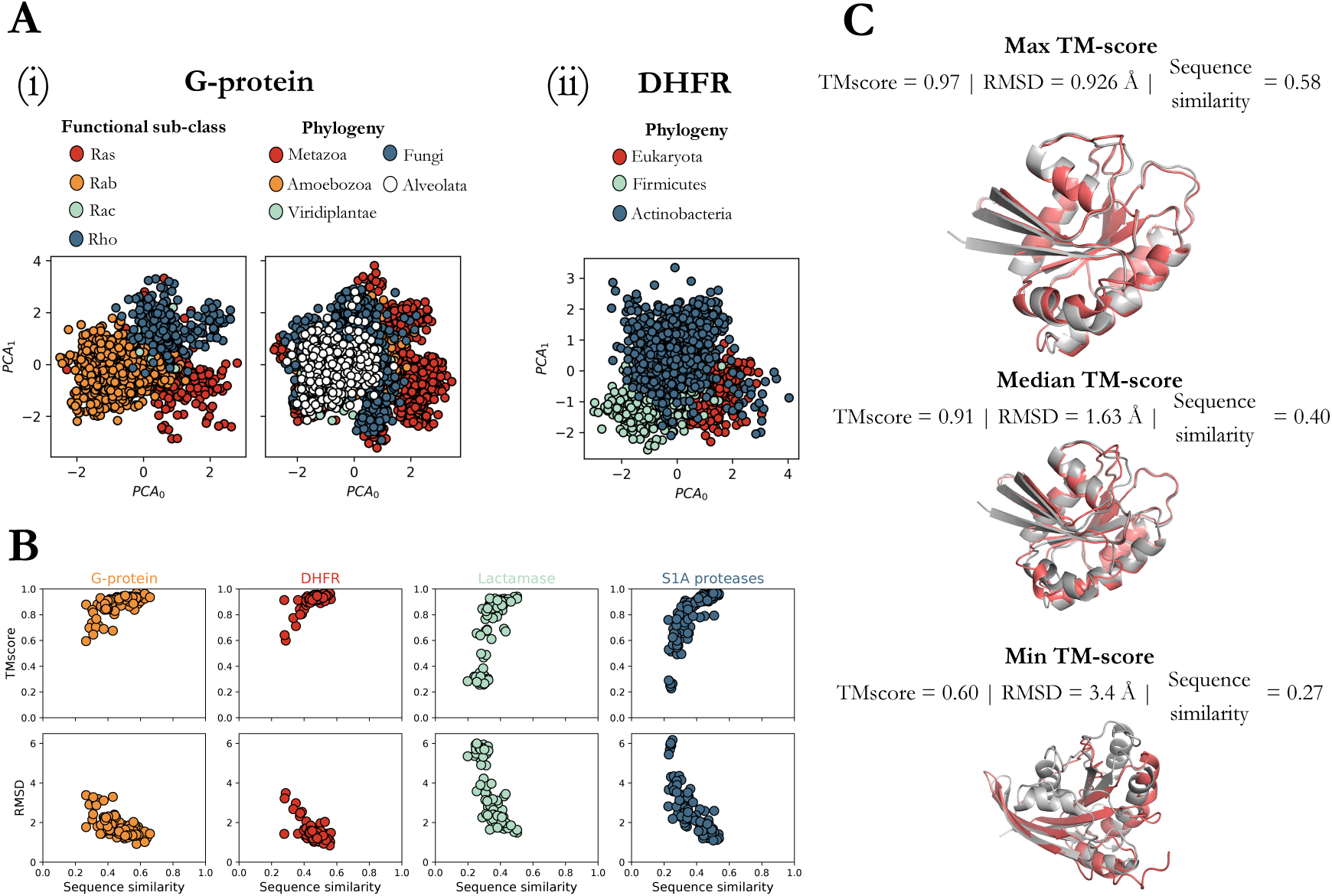
ProtWave-VAE can infer meaningful biological representations on alignment-free protein families. (A) Here, we present the principal component analysis (PCA) projections of the inferred latent spaces for the (i) G-protein and (ii) DHFR families. For the G-protein family, we find that the unsupervised model disentangles homologs within the inferred latent space based on functional sub-classes and phylogeny. Similarly, with the DHFR family, the model learns to disentangle homologs in the inferred space in terms of the phylogeny. (B) To test ProtWave-VAE generative capacity, we randomly sampled 100 latent vectors *z* for each protein family from a normal distribution *N* (0,I), corresponding to the latent prior. Then using a computational structure prediction workflow (ColabFold + TMalign), we predicted each structure of the sample sequences and compared the predicted structure against a natural homolog that defines the corresponding protein family, retrieving TMscores and root-mean-square distance (RMSD) scores. We computed the minimum Levenshtein distance between the sampled novel sequence and training sequences normalized the length of the longer sequence in the pair. (C) Using G-protein structure predictions of ProtWave-VAE novel design sequences (red), we visualize the alignment of maximum, median, and minimum TM-score synthetic sequences (grey). Figure S2 illustrates the latent spaces, structure predictions, and TMscores of the remaining protein families.

Our computational results suggest that combining an autoregressive decoder with a latent-based inference model permits inference of meaningful biological representations within a learned low-dimensional latent space. However, can our approach still generate novel samples indistinguishable from the training data distribution? To test the generated alignment-free sequences, we used AlphaFold2 with MMSeqs2 (i.e., ColabFold^31,32^) to predict the tertiary structure of the generated sequences and determine whether the predictions recapitulate the tertiary structure of the natural homologs. To showcase our model’s ability to infer meaningful representations from protein families, we randomly sampled 100 latent vectors from an isotropic Gaussian distribution and used the trained ProtWave-VAE WaveNet-based autoregressive decoder to generate novel sequences for each protein family. To measure the novelty of the generated sequences, we computed the minimum Levenshtein distance from any homolog in the training dataset normalized by the length of the longer sequence within the pair. Additionally, we sought to assess whether the generated sequences were predicted to adopt a tertiary fold consistent with the homologous family. We compared the predicted tertiary structure of each ProtWave-VAE designed sequence to a prototypical member of the homologous family with a known crystal structure available within the Protein Data Bank: ^39^ 5P21 for G-protein, 1XR2 for DHFR, 3TGI for S1A serine protease, and 1FQG for beta-lactamase. To quantify the similarity of the folds, we computed the TMscore and heavy-atom root-mean-squared distances (RMSD) using the TMalign algorithm.^40^ We present scatter-plots of the TMscore against sequence novelty and RMSD against sequence novelty for the 100 designed proteins in Figure 2B. Our results show that the artificial proteins possess TMscores in the range of 0.2-1.0^41^ and RMSDs in the range of 0-6 Å, with TMscores having a strong positive correlation with sequence similarity and RMSDs having a strong negative correlation with sequence similarity.

Furthermore, we demonstrate the model’s ability to generate sequences with tertiary structures similar to the native fold without being exposed to any structural information during training. In Figure 2C and Figure S2B, we show the AlphaFold predicted tertiary structures of three representative synthetic sequences with maximum, median, and minimum TM-scores. The similarity of the synthetic sequence tertiary structures to the native folds as quantified by high TMscores and low RMSDs for all four protein families supports that the correlated patterns of amino acid mutations learned by the model within unaligned sequence data are sufficient to generate tertiary structures representative of the homologous family’s native fold. In summary, the ProtWave-VAE model demonstrates the capability to infer meaningful biological representations and generate novel sequences with tertiary structures with native-like folds.

### Fitness landscape benchmarking using semi-supervised learning

We subsequently evaluated the capabilities of the ProtWave-VAE model in semi-supervised downstream fitness prediction tasks and compared its performance against a number of leading methodologies. The primary rationale behind semi-supervised learning approaches is that latent representations *z* become more informative for predicting downstream functional properties *y* when they are also employed for reconstructing sequences *x*.^33,42^ In a similar vein, it is plausible to suggest that unlabeled protein sequences contain considerable information about their structure and function.^43,44^ In addition, semi-supervised learning is beneficial when labels are scarce, and unlabeled data is abundant, which is generally the case for protein design applications where only a small fraction of entries within large sequence databases are annotated with functional assays. One advantage of the AR-VAE architecture of ProtWave-VAE is that it can employ semi-supervised learning via its learned latent space in a straightforward manner that is difficult to achieve for standalone AR approaches. We benchmark our model on four popular semi-supervised downstream function and fitness prediction tasks from two popular community benchmarks: Task Assessing Protein Embeddings (TAPE)^35^ and Fitness Landscape Inference for Proteins (FLIP)^34^ baselines. The four semi-supervised prediction tasks are: (1) the highly epistatic mutational landscape of GB1 (FLIP),^45^ (2) mutational screening of the fitness landscape of VP-1 AAV (FLIP),^46,47^ (3) stability landscape prediction (TAPE),^48^ and (4) epistatic green fluorescent protein (GFP) landscape prediction (TAPE).^49^

Our benchmark evaluations show that ProtWave-VAE either rivals or surpasses a number of state-of-the-art (SOTA) models, including large language models – ESM-1b and ESM-1v –^50,51^ transformer-based models – TAPE Transformer^35,52^ and masked dilated convolution-based architectures – CARP-640M^53^ (Table 1). In the GB1 task, ProtWave-VAE exceeds SOTA models in random split and 3-vs-rest split, while remaining competitive in the 1-vs-rest and 2-vs-rest split based on the Spearman *ρ* rank correlation. These findings imply strong extrapolation capabilities in genotype space for the model. Nonetheless, it underperforms on the low-vs-high split in the GB1 dataset, suggesting effective learning from low fitness mode (training samples) but more limited accuracy in extrapolating and capturing high fitness mode (test samples). In the AAV task, where the model aims to predict fitness measurements of AAV capsid mutants, ProtWave-VAE surpasses SOTA models in random splits, design-mutant split, and 1-vs-rest, while competing effectively on mutant-design split and 2-vs-rest against large language models using either transformer or dilated convolution architectures. Only in the stability task does our model underperform in stability regression predictions based on the Spearman *ρ* correlation, and again remains on par with SOTA models when it comes to fluorescence regression prediction in the GFP task.

**Table 1:**
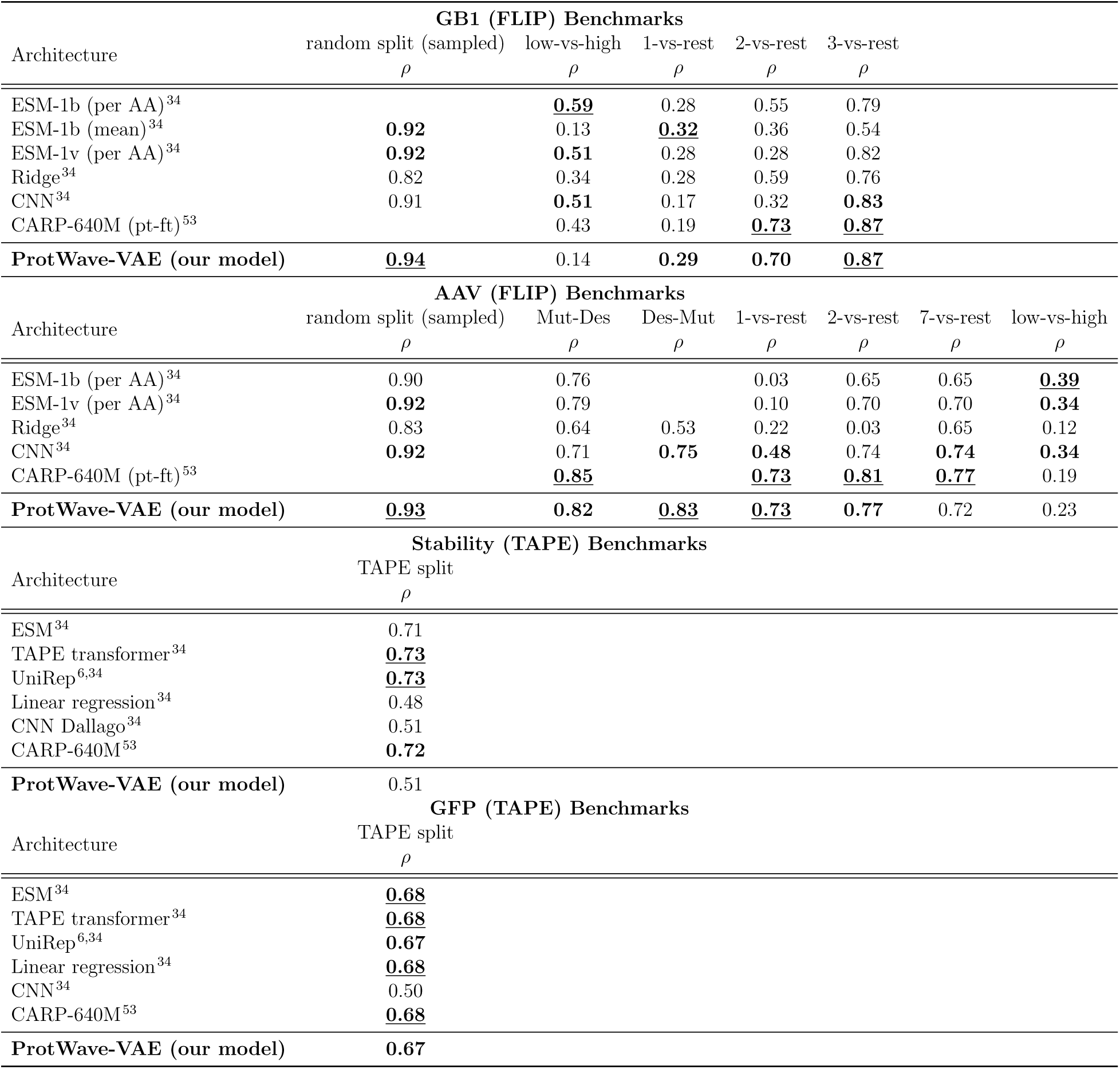
Protein property prediction results on FLIP and TAPE benchmark. The **<u>best</u>** results and the **second-best** results are marked.

Overall, these results demonstrate that the ProtWave-VAE model performs well on semi-supervised downstream functional and fitness prediction at a level competitive with state-of-the-art models such as ESM and CARP-640M. This demonstrates that ProtWave-VAE is learning internal latent space representations that expose the ancestral and functional relationships necessary to both generatively design novel synthetic sequences and accurately predict their functional properties. Furthermore, we observe that the ProtWave-VAE is much more lightweight than typical SOTA models, containing 100-fold fewer trainable parameters that the lightest-weight transformer model (ESM^50,51^) considered in the benchmark suite, conveying advantages and savings in cost and time for model training and deployment.

### Generative modeling with semi-supervised learning for the Chorismate mutase family

We next sought to explore the potential impact of incorporating experimental knowledge into generative learning tasks by employing semi-supervised learning to reshape the latent space based on functional measurements.^54^ The goal of many protein engineering campaigns is to design proteins with elevated function along one or more functional dimensions. We hypothesized that the incorporation of functional measurements into training of the ProtWave-VAE within a semi-supervised paradigm could induce functional gradients within the learned latent space and partially disentangle the latent representation to foreground the functional property of interest. We further hypothesized that this reshaped latent space would support superior conditioning and generative decoding of synthetic mutants with elevated function. To test these hypotheses, we compared unsupervised and semi-supervised learning for the Chorismate mutase (CM) protein family. ^36^ We found that semi-supervised learning infers a gradient in fitness whereas unsupervised learning does not (Figure 3A), indicating that information from experimental assays can be leveraged to sculpt the latent space to induce gradients in properties of interest. To test whether the introduction of the second decoder for the semi-supervised regression task harm the model’s generative capacity, we generatively designed 100 sequences for both unsupervised and semi-supervised trained models and computed the TMscore and RMSD scores between predicted design structures and natural *Escherichia coli* crystal structural (PDB:1ECM) using ColabFold structure predictions (Figure 3B,C). Our results demonstrate no performance degradation in terms of TMscore or RMSD values between unsupervised and semi-supervised and show that the predicted design structures accurately recapitulate the native fold. This result demonstrates that semi-supervised learning can reshape the latent space to induce a gradient in a property of interest while maintaining the generative capabilities of the ProtWave-VAE model. Determining whether the designed synthetic CM sequences do indeed possess elevated function as we advance up, and extrapolate beyond, the induced functional gradient can, of course, only be ascertained by experimental gene synthesis and functional assays.

**Figure 3:**
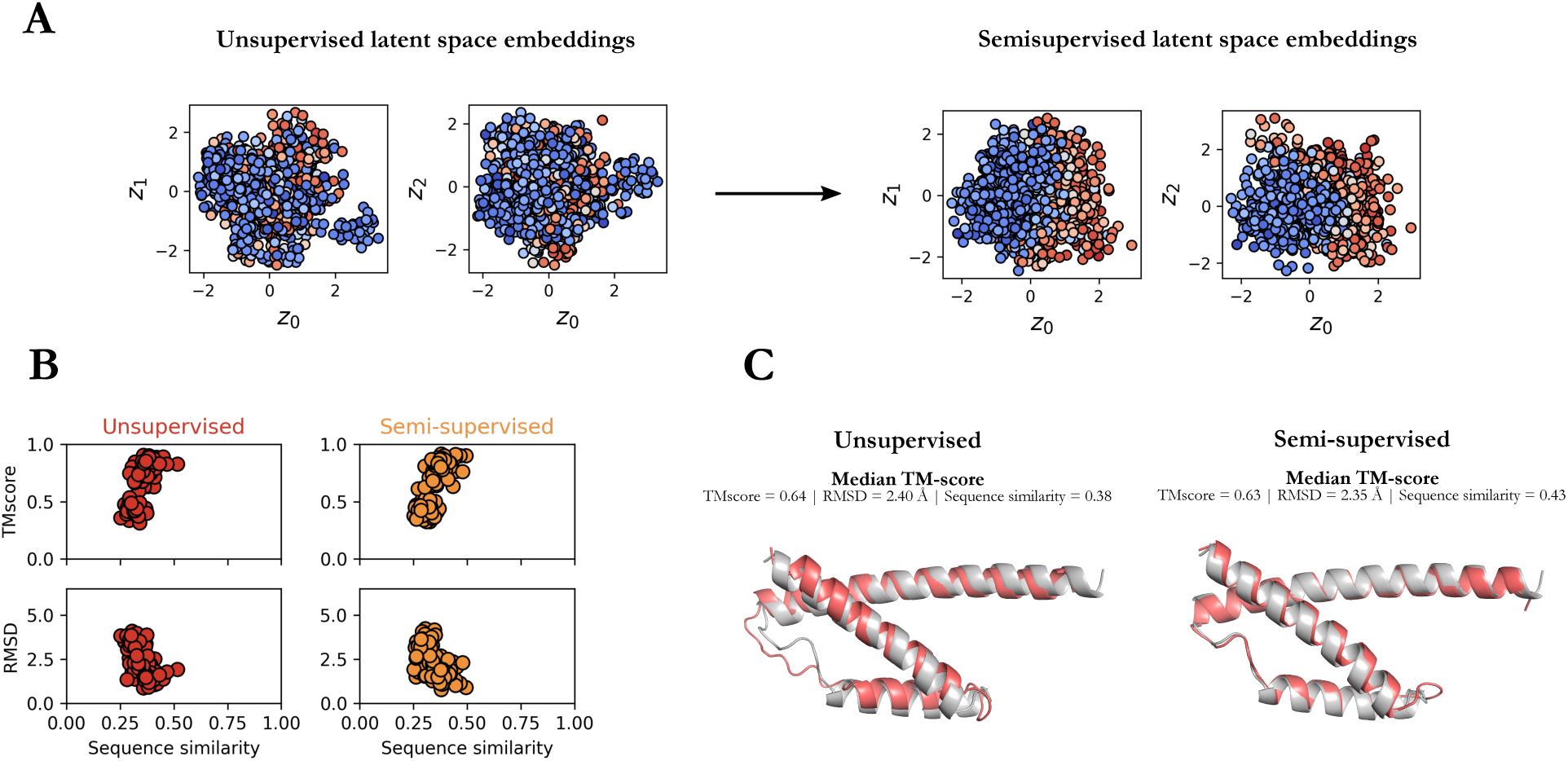
Comparison of unsupervised and semi-supervised learning for the generative design of the Chorismate mutase (CM) proteins. (A) Semi-supervised learning allows us to infer and reshape the latent space so that latent coordinates correlate more strongly with CM fitness as measured by relative enrichment select-seq assay scores. (B) To verify that reshaping the latent space does not lead to a loss of generative capabilities for the model, we used the ColabFold plus TMalign algorithm to demonstrate no significant loss of generative performance in the RMSD and TMscores of the predicted tertiary structures generated by unsupervised and semi-supervised ProtWave-VAE models. (C) A superposition of the wild-type *E. coli* crystal structural (PDB:1ECM) (grey) and the ColabFold predicted structure of the ProtWave-VAE designed sequence possessing the median TM score (red), shows excellent agreement for both the unsupervised and semi-supervised models.

### Introducing novel latent-based autoregression for sequence diversification

The integrated AR-VAE architecture of the ProtWave-VAE model enables a potentially useful form of synthetic protein generative design that we refer to as C-terminus diversification with latent conditioning. By combining latent inference, conditioning, and autoregressive amino acid generation, our model allows us to condition on both the latent vector within the VAE latent space and an arbitrary number of N-terminal residues in the generated protein to diversify the C-terminal region. Simple latent-based generative models such as standard VAEs allow for generating a whole sequence by conditioning on latent embeddings, but typically cannot also condition on amino acids from a known natural protein of interest and then diversify (i.e., “inpaint”) the missing region of interest. In contrast, AR generative models generate sequences by predicting subsequent amino acid while conditioning on previously predicted amino acids, but cannot conduct latent inference and use those latent embeddings to control the design of synthetic sequences. Autoregressive generation in this manner has proven to be successful strategy and valuable tool in, for example, nanobody design, by diversifying the complementary determining region CDR3 while conditioning on CDR1 and CDR2.^7^ However, the absence of a biologically meaningful latent space to condition latent codes to inpaint missing regions means that the generation process cannot be readily guided to introduce particular ancestral or functional characteristics. We propose that the capability of the ProtWave-VAE to perform simultaneous N-terminal and latent conditioning may prove valuable in applications to nanobodies, antibodies, enzymes, signaling domains, linkers, and multimeric proteins, where it is desirable to maintain some structural and/or functional properties of the N-terminal region and introduce new capabilities by re-design of the C-terminus. We demonstrate this novel generative approach in an application to C-terminal diversification of the *E. coli* CM homolog (PDB: 1ECM).

We sampled 100 latent embeddings from a normal distribution *N* (0,I) over the latent space of the unsupervised / semi-supervised ProtWave-VAE (Figure 3). These latent codes were then used to perform latent-only conditional synthetic generation of 100 novel CM sequences (Algorithm 1). We then performed N-terminus plus latent conditional generation of an additional 100 CM sequences using the same latent codes, but where we also fixed the identity of the N-terminal residues 1-40 and inpainted the remaining C-terminal residues 41-96 using autoregressive sampling (Algorithm 2) (Figure 4A). We hypothesized that the sequences generated by the two approaches should possess ColabFold predicted tertiary structures in equally good agreement to the wild-type *E. coli* crystal structure, but that the C-terminal region (residues 41-96) should follow a different distribution in the two ensembles reflecting the impact of N-terminal conditioning on the autoregressive generation process. Specifically, the sequences generated by latent-only conditioning should access a more diverse sequence space compared to those additionally constrained by N-terminal conditioning on the wild-type sequence.

**Figure 4:**
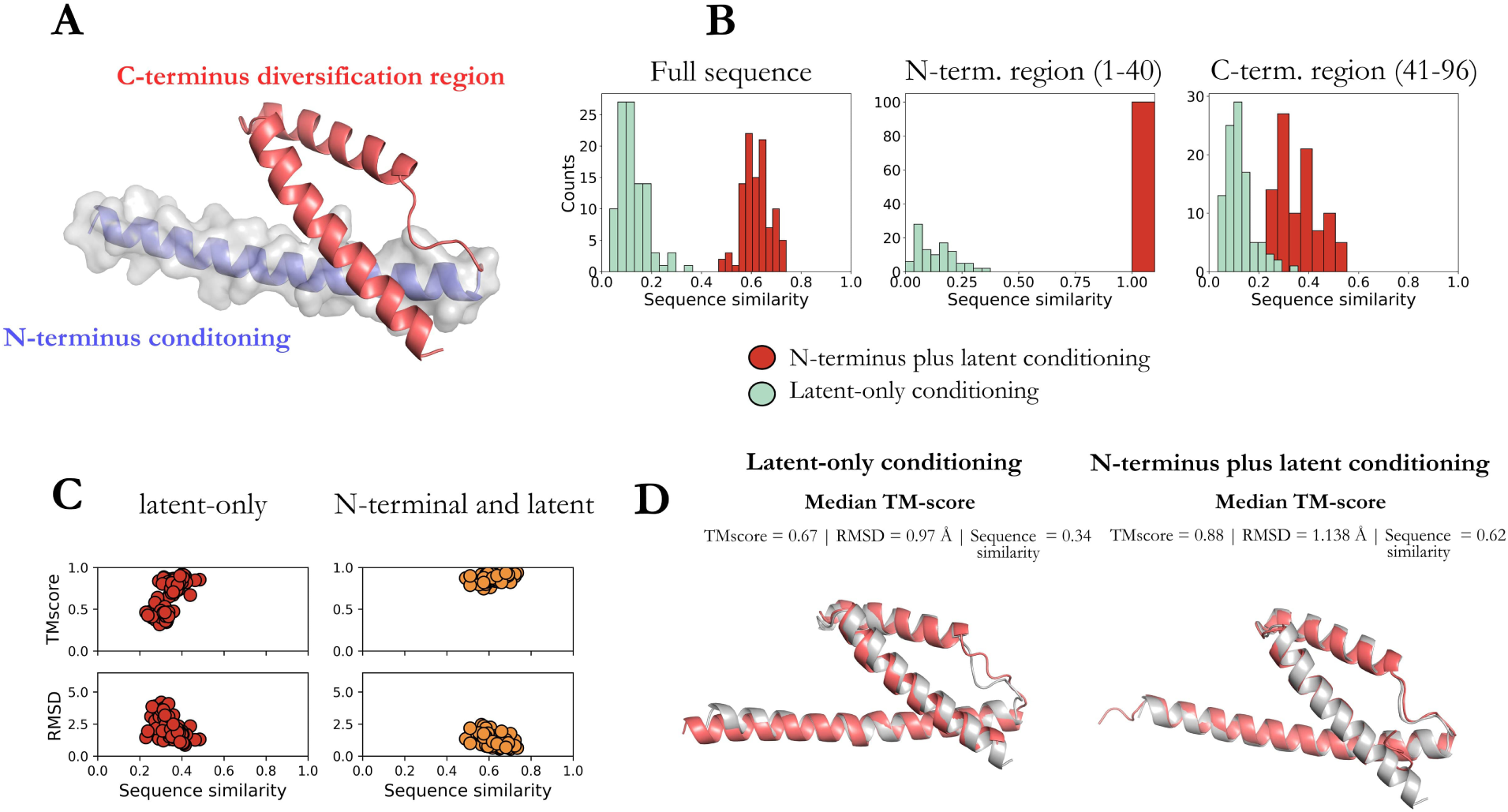
Introduction of C-terminus diversification with N-terminus and latent conditioning. (A) Tertiary structure of the E. coli CM wild-type protein (PDB: 1ECM) illustrating the N-terminal region (residues 1-40) used for N-terminal conditioning (blue) and the remaining C-terminal region (residues 41-96) subject to generative diversification (red). (B) We generated 100 novel latent conditioning vectors by sampling from a *N* (0,I) prior over the latent space and used each latent vector to generate a novel synthetic protein with and without N-terminal conditioning. We compare the sequence similarity of the full sequence, N-terminal conditioning, and C-terminal diversification regions between the latent-only (green) and N-terminal plus latent (red) conditioned generated sequence ensembles.(C) ColabFold predicted tertiary structures show no significant differences in the RMSD and TMscores between the latent-only and N-terminal conditioned designs. (D) Structure prediction (red) of the median TMscore sequence for both latent-only and N-terminal plus latent conditioned designs against the *E.coli* wild-type crystal structure PDB:1ECM (grey).

#### Algorithm 1 Latent-only conditioning

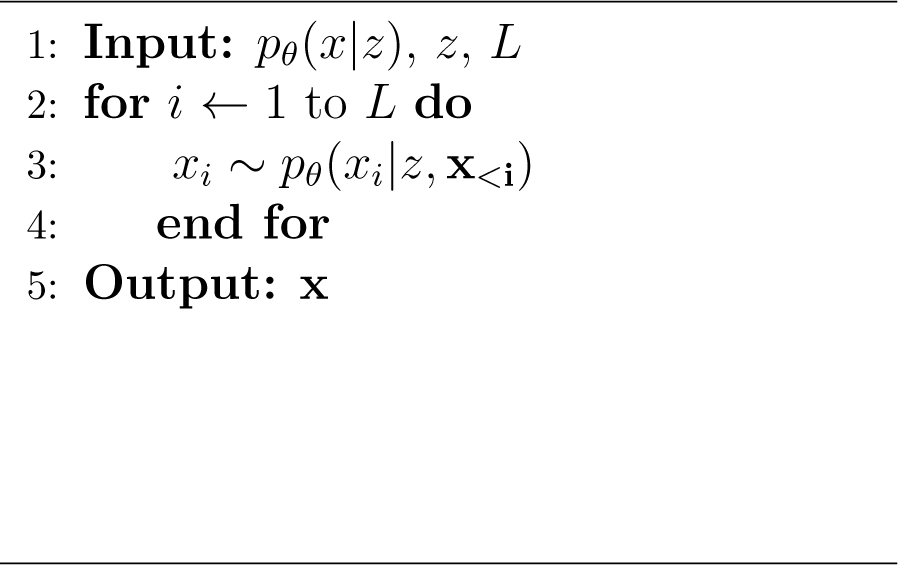

#### Algorithm 2 N-terminus plus latent conditioning

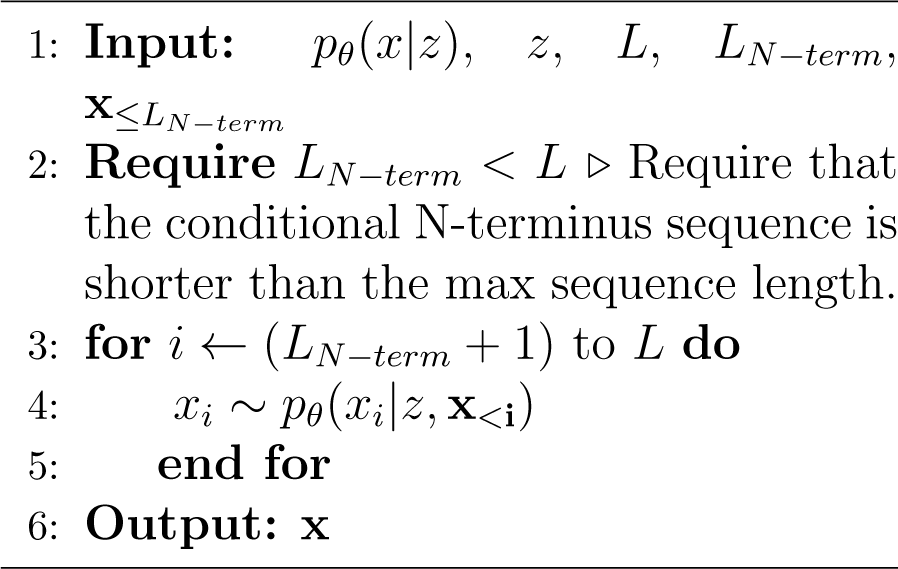

As anticipated, the sequences designed by latent-only conditioning were more dissimilar to the *E. coli* wild-type than those produced by N-terminal and latent conditioning (Figure 4B). Of course, this follows because the N-terminal region comprising residues 1-40 is identical to the wild-type for all sequences generated by the N-terminal conditioned approach, resulting in 100% sequence similarity within the N-terminal region for these sequences, whereas the latent-only sequences possess sequence similarities in the range of 0-50%. Further, the sequence similarity of the C-terminal region comprising residues 41-96 is higher for the N-terminal and latent conditioned sequences than the latent-only sequences. This is a direct result of the autoregressive nature of the model wherein the fixed N-terminal region conditions the generation of the C-terminal region to remain closer to the wild-type for the same latent vector. ColabFold structure predictions and RMSD and TMscore evaluation of the generated protein sequences demonstrate the anticipated positive correlation between sequence similarity and TMscore and negative correlation between sequence similarity and RMSD (Figure 4C). The smaller diversity of the N-terminal and latent conditioned sequences mean that they exhibit a tighter distribution than those produced by latent-only conditioning. Comparison of the ColabFold structure predictions for the synthetic sequences with the median TMscore show them to possess tertiary structures visually indistinguishable from the *E. coli* wild-type crystal structure for both the N-terminal and latent conditioning and latent-only conditioning generation processes (Figure 4D).

Taken together, these results demonstrate that N-terminal conditioning can be used to effectively constrain the degree of C-terminal diversification by providing additional conditioning of the autoregressive sequence generation process. The structural similarity of the synthetic sequences to the native fold of the protein family, at least for this application, is insensitive to whether or not N-terminal conditioning is used in the generation procedure.

### Experimental validation of latent-based autoregression for *de novo* **design and natural sequence diversification**

So far, we have demonstrated the ability of the ProtWave-VAE model to infer biologically meaningful latent space embeddings, perform downstream functional prediction tasks competitive with state-of-the-art approaches, operate in a semi-supervised fashion without compromising generative capacity, and perform N-terminal conditioned sequence generation. All of these demonstrations have been performed by retrospective analysis of existing data sets and comparison of tertiary structures generated by ColabFold. To rigorously validate our ProtWave-VAE capabilities in functional protein design, it is necessary to experimentally synthesize and assay the sequences generatively designed by the model. To do so, we trained a ProtWave-VAE model to design synthetic Src homology 3 (SH3) sequences capable of functioning like natural SH3*^Sho^*^1^ domains by binding its cognate pbs2 ligand and effecting the osmosensing mechanism in *Saccharomyces cerevisiae* as assessed by a a select-seq assay^14,37^ (Figure 5A). This assay couples a high-osmolarity challenge with next-generation sequencing to measure the relative enrichment (r.e.) of the post-selection population in a particular mutant relative to a null gene and the *S. cerevisiae* wild-type.^14^ The r.e. score provides a quantitative measurement of the degree to which our designed SH3 domains are functional *in vivo* and capable of activating a homeostatic osmoprotective response. The assay shows good reproducibility in independent trials (*R*^2^ = 0.94; Figure S3). We trained a semi-supervised ProtWave-VAE model on an SH3 dataset consisting of natural SH3 proteins mostly from the fungal kingdom and synthetic proteins designed using our previous generation unsupervised learning VAE models for which we possess functional assay measurements.^14^ The semi-supervised nature of the model is observed to induce a strong r.e. gradient in the latent space (Figure 5B), allowing us to condition generative sequence design by drawing latent vectors in the functional region of the latent space using our latent-only (Algorithm 1) and N-terminal and latent conditioned (Algorithm 2) strategies (Figure 5C).

**Figure 5:**
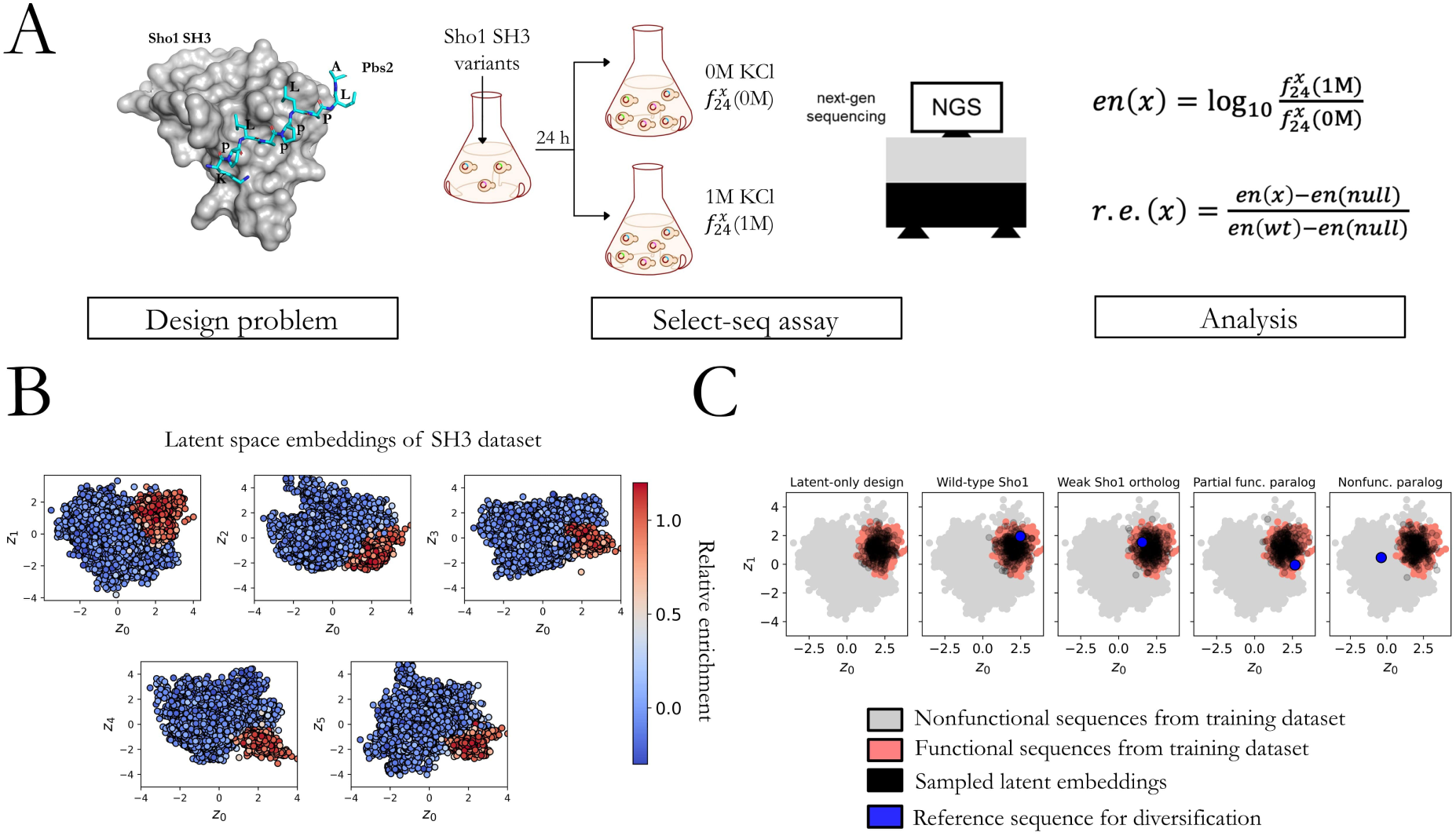
Experimental assessment of ProtWave-VAE generatively-designed synthetic Sho1*^SH^*^3^ domains. (A) The crystal structure (PDB: 2VKN) of the *S. Cerevisiae* wild-type (wt) Sho1*^SH^*^3^ binding to the pbs2 ligand (blue sticks) is shown along with an illustrative cartoon of the select-seq assay and next-gen sequencing platform for measuring relative enrichment (r.e.) scores as a measure of fitness. The enrichment *en*(*x*) of mutant protein *x* is the logarithm of the ratio *f ^x^* (1*M*)/*f ^x^* (0*M*), where *f ^x^* (*kM*) is the frequency of mutant *x* in the population after being subjected to 24 h of *k*M KCl solution. The relative enrichment *r.e.*(*x*) of mutant *x* is a normalized enrichment score relative to the wild-type protein and a null protein such that *r.e.*(*x*)=1 indicates the same functional performance as the wildtype and *r.e.*(*x*)=0 indicates the same functional performance as the null gene. (B) The six-dimensional latent space spanned by latent vectors **z** = *{z*_0_*, z*_1_*, z*_2_*, z*_3_*, z*_4_*, z*_5_*}* of a trained semi-supervised ProtWave-VAE model exposes clear gradients in the r.e. scores in all 2D projections of this space. The high fitness training sequences (red) are clustered and segregated from the low-fitness training sequences (blue). (C) We generated synthetic ProtWave-VAE sequences for experimental testing by five separate protocols: (I) latent-only conditioning, (II) N-terminal and latent conditioning of wild-type Sho1*^SH^*^3^, (III) N-terminal and latent conditioning of a weak binding Sho1 ortholog, (IV) N-terminal and latent conditioning of a partial rescuing SH3 paralog, and (V) N-terminal and latent conditioning of a SH3 paralog that does not rescue Sho1 functionality. We illustrate the nonfunctional (grey, r.e.*<*0.5) and functional (red, r.e.*≥*0.5) sequences within the *z*_0_-*z*_1_ projection of the latent space together with the sampled latent space embeddings (black) and, if appropriate, reference sequence used for N-terminal conditioning (blue). In all cases, latent vectors were drawn from the region of the latent space containing the functional training sequences so as to guide the generation of functional synthetic Sho1*^SH^*^3^ orthologs. We did so by fitting an anisotropic Gaussian to the r.e.*≥*0.5 (red) training points and randomly sampling from this distribution to generate the latent conditioning vectors (black).

We employed the trained ProtWave-VAE model to devise sequences using five distinct protocols, resulting in five subgroups of designed sequences (I-V). Subgroup (I) utilized latent vectors for conditioning the *de novo* generative design of synthetic sequences. By fitting a 6D anisotropic Gaussian to the embedding of training sequences with select-seq measurements of r.e.*≥*0.5 and randomly sampling latent vectors from this distribution (Figure 5C, black points), the latent vectors were extracted from the ProtWave-VAE latent space. This ensured the latent vectors originated from a region of latent space containing functional training sequences, which in turn conditioned the generation of functional sequences. The other four subgroups were designed using the N-terminal and latent conditioning approach. To assess our model’s capability in producing functional designs, we chose four reference proteins for N-terminus conditioning, defining subgroups (II) wild-type *S. cerevisiae* Sho1*^SH^*^3^, (III) a weak binding Sho1 ortholog, (IV) a partial rescuing SH3 paralog, and (V) a nonfunctional *S. cerevisiae* paralog. The latent embeddings for these conditioning sequences are denoted by blue points in Figure 5C. We observed that the nonfunctional SH3 paralog was situated outside the functional niche, as it could not rescue Sho1 functionality. In the latent-only *de novo* design subgroup (I), we generated 150 sequences. In the four N-terminal and latent conditioned subgroups (II-V), we produced 150 (II-IV) and 99 (V) designs subdivided into three equal-sized sub-categories differentiated by the fraction of the sequence used for N-terminal conditioning: 25%, 50%, and 75%. Subgroups (II-V) were intended to assess various protein design goals, including: (II) preserving function, (III-IV) enhancing existing function, and (V) gain of function. Finally, we included two control subgroups (VI) and (VII), each comprising 150 sequences, for the N-terminal and latent conditioning subgroups (III) – N-terminal conditioning on a weak binding Sho1 ortholog – and (IV) – N-terminal conditioning on a partial rescuing SH3 paralog – to verify that our model’s capacity to improve function was not simply due to chance. Within these control subgroups, we introduced random mutations in the designable C-terminal region until the distribution of the 50 sequences within each subgroup sub-category matched the Levenshtein distances to the *S. cerevisiae* wild-type of the 50 sequences designed by the ProtWave-VAE (Figure S1).

Figure 6A displays the experimental outcomes for sequences belonging to the latent-only conditioning subgroup (I). Out of the 150 gene designs, 148 were successfully assembled and tested experimentally. The first two plots are scatterplots, where each point represents a designed sequence, the *y*-axis denotes the relative enrichment score, and the *x*-axis indicates the sequence similarity relative to any sequence in the training dataset and wild-type SH3*^Sho^*^1^ *S. cerevisiae*. For the latent-only conditioning subgroup (I), the designed sequences exhibit the ability to rescue functionality (i.e., r.e.*≥*0.5) while covering a broad spectrum of normalized Levenshtein distances, ranging from sequence similarities of 75-100% and 45-70% relative to the training dataset and wild-type SH3*^Sho^*^1^. This suggests that the ProtWave-VAE model can produce highly diverse sequences that significantly deviate from the wild-type, yet retain the correlated amino acid residue patterns necessary for maintaining osmosensing function. The final bar graph highlights the superior performance of ProtWave-VAE compared to our previously reported InfoVAE and VanillaVAE models when designing sequences using local sampling.^14^ The ProtWave-VAE outperforms the two VAE models, rescuing functionality at a level of 51% (76 out of 148) versus 48% and 21%, respectively. In addition to its high performance in designing *de novo* functional sequences with latent-only conditioning, ProtWave-VAE can, unlike our previous two VAE approaches, does not require training over a multiple-sequence alignment and can readily generate variable-length sequences.

**Figure 6:**
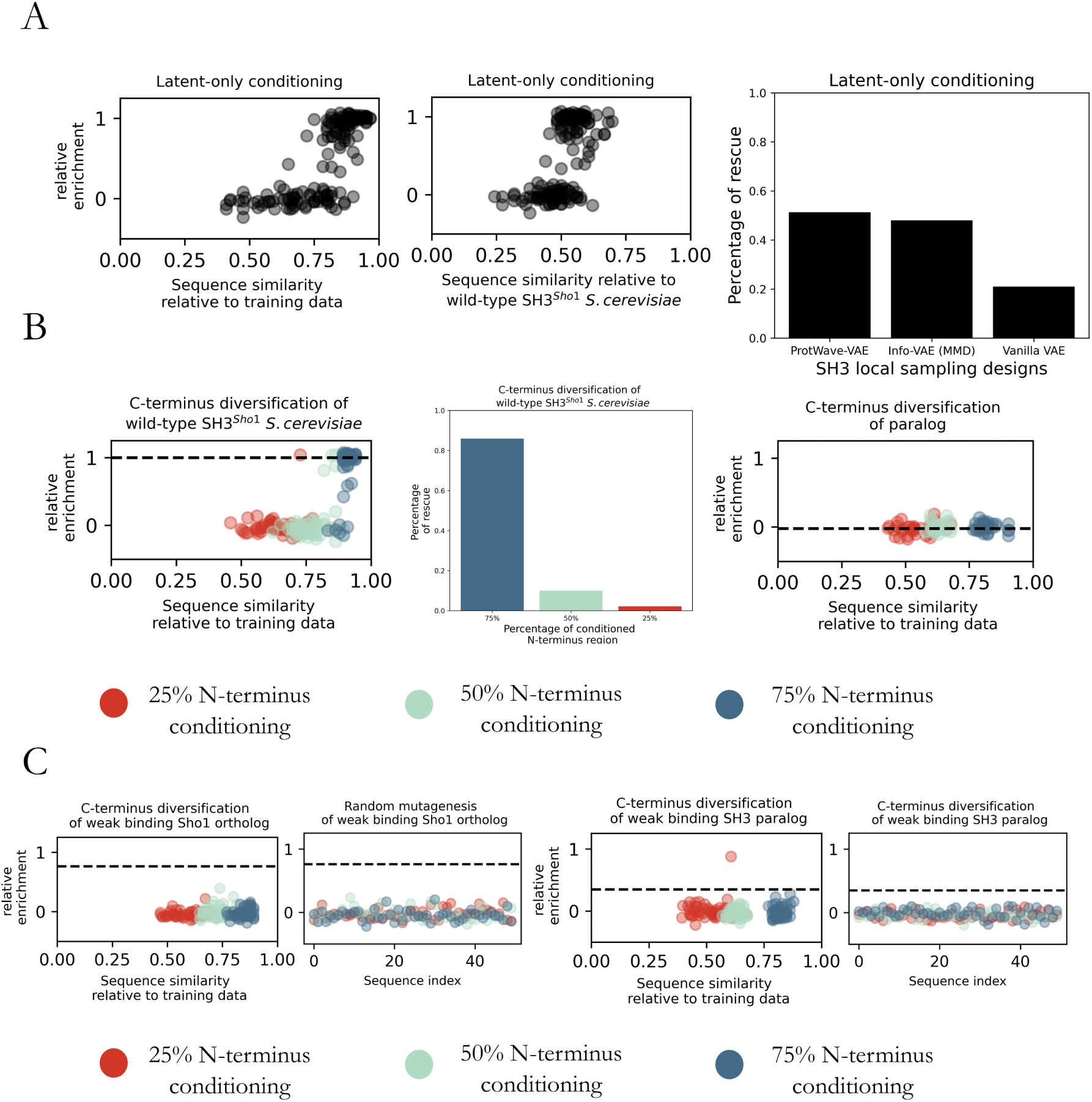
Experimental outcomes of ProtWave-VAE generated sequences. (A) Subgroup (I) – latent-only *de novo* design. Scatterplots illustrate the sequence similarity measured by normalized Levenshtein distance and relative enrichment (r.e.) scores for the synthetic designs. The bar graph demonstrates ProtWave-VAE performance through local sampling compared to the synthetic generative designs previously reported for our VAE-based DGM using an Info-VAE employing a max-mean discrepancy (MMD) loss and a Vanilla VAE employing the standard ELBO loss.^14^ The ProtWave-VAE generates diverse sequences with a high probability of functional rescue. (B) Subgroups (II) – maintaining function with C-terminus diversification for SH3 wild-type – and (V) – elevating function of a nonfunctional paralog using C-terminus diversification. Experimental measurements for design groups employing 25% (red), 50% (green), and 75% (blue) of the sequence length for N-terminal conditioning. The scatterplot on the left displays the r.e. scores for subgroup (II) versus sequence similarity to the training dataset. The bar graph reveals the rescue percentage within the design pool for subgroup (II) at varying N-terminus conditioned percentages. The scatterplot on the right presents r.e. versus sequence similarity to the training dataset for subgroup (V). None of these sequences rescued osmosensing function. The horizontal dotted line for both scatterplots corresponds to the relative enrichment score of the homolog used for N-terminus conditioning. (C) Subgroups (III) and (VI) – elevating function of a weak binding Sho1 ortholog using C-terminus diversification and its random mutagenized control – and (IV) and (VII) – elevating function of a partial rescuing SH3 paralog using C-terminus diversification and its random mutagenized control. The two left scatterplots pertaining to subgroups (III) and (VI) failed to show any rescue at any level of N-terminal conditioning. The two right scatterplots pertaining to subgroups (IV) and (VII) show that one of the generatively designed sequences with 25% N-terminal conditioning did rescue function.

Figure 6B showcases the experimental outcomes for sequences belonging to the N-terminus plus latent conditioning for C-terminus diversification of the wild-type SH3*^Sho^*^1^ – subgroup (II) – and paralog SH3 that fails to rescue function – subgroup (V). Considering first the left scatterplot displaying the r.e. measurements and sequence similarities for the generatively designed sequences in subgroup (II) that were successfully assembled and tested, we find the number of functional sequences for each N-terminus percentage prompt to be 1 out of 46 for 25% N-terminal conditioning (red), 5 out of 50 for 50% (green), and 43 out of 50 for 75% (blue). The bar graph presents the percentage of rescue, which amounts to 86%, 10%, and 2.2% for 75%, 50%, and 25% N-terminal conditioning, respectively. These results suggest that conditioning on both the N-terminus and the latent vectors to inpaint the remaining C-terminus can lead to functional synthetic sequences, but that the success rates decrease with increasing degree of C-terminus inpainting and number of amino acid positions to diversify. We propose that this decaying success rate may result from an incompatibility of the two conditioning goals such that the latent vector seeks to generate a C-terminal sequence incompatible with the pre-defined N-terminal sequence. Turning to the right scatterplot presenting the data for the the C-terminus diversified paralog designs in subgroup (V), we find that none of the generatively designed sequences that were successfully assembled and subjected to experimental testing were capable of rescuing function: 0 out of 32 for 25% N-terminal conditioning (red), 0 out of 33 for 50% (green), and 0 out of 33 for 75% (blue). Together with the potential incompatibility of the two conditioning goals, it is also possible that if any N-terminal residues are indispensible to protein function – either directly through binding to the target ligand or indirectly via their participation in critical multi-body interaction networks with other amino acids – and these are conditioned to contain mutant residues via the N-terminal conditioning, then no C-terminal inpainting can lead to functional rescue.

Figure 6C displays the experimental outcomes for sequences belonging to the N-terminus plus latent conditioning for C-terminus diversification of the weak ortholog SH3*^Sho^*^1^ – sub-group (III) – and partial paralog SH3 – subgroup (IV). The design goal for these two subgroups was to enhance functionality by inpainting the C-terminus. As a control to demonstrate that the generative model performs better than random mutagenesis, we experimentally tested mutants in subgroup (VI) and (VII). The first scatterplot corresponds to sub-group (III). Similar to our results for the SH3 paralog – subgroup (V) – we find that none of the designed sequences were able to rescue function: 0 out of 50 for 25% N-terminal conditioning (red), 0 out of 50 for 50% (green), and 0 out of 50 for 75% (blue). The second scatterplot contains data for the subgroup (VI) control, in which we randomly mutated the C-terminal region of the weak ortholog to produce the same distribution of Levenshtein distances to the *S. cerevisiae* wild-type as generated in the (III) treatment group. Again, we observe no sequences capable of functional rescue: 0 out of 50 for 25% N-terminal conditioning (red), 0 out of 50 for 50% (green), and 0 out of 50 for 75% (blue). These results indicate that ProtWave-VAE was unable to improve functionality of the weak ortholog using N-terminus and latent conditioning.

The third scatterplot in Figure 6C corresponds to subgroup (IV), in which we considered C-terminal diversification of a partial rescuing SH3 paralog. In this case we observe one success out of the 48 designed sequences with 25% N-terminal conditioning (red), that resulted in a boost of the r.e. score from an initial value of 0.35 for the partially rescuing paralog to a value of 0.88, corresponding to a 2.5*×* enhancement in functionality by inpainting the missing C-terminus region. Interestingly, with a sequence similarity of just 61% to any training sequence, this C-terminus diversified design is the most novel functional artificial sequence even when compared to any of the *de novo* designs with latent-only conditioning in subgroup (I). None of the other generated sequences – 0 out of 50 for 50% (green), and 0 out of 50 for 75% (blue) – were capable of rescue. The control subgroup (VII) is presented in the fourth scatterplot, within which no sequences exhibited functionality. This result implies that N-terminus plus latent-only conditioning can serve as an approach for designing novel sequences by conditioning on distant paralogs and inpainting the C-terminus to elevate function, although the overall hit rate of functional sequences is relatively low. Again, we conjecture that this may result from an inherent incompatibility of the latent vector and N-terminal conditions, and that the successful rescue resulted from a generative design in which these conditions were mutually compatible.

The experimental results reveal that our proposed model can effectively design synthetic proteins with functional properties on par with their natural counterparts. Additionally, the model is capable of venturing into unexplored regions of sequence space that have not been traversed by natural evolution. Our study also presents a novel protein engineering technique for diversifying the C-terminus of proteins, contributing to the preservation and enhancement of their functionality. Among the findings, the most noteworthy is the ability of the ProtWave-VAE model to imbue a weak binding SH3 paralog from the Hof1 paralog group with osmosensing function, resulting in a 2.5*×* increase in relative enrichment and generating the most novel sequence among all synthetic designs sharing only 61% sequence similarity to any training sequence.

## Conclusion

In this work, we introduce ProtWave-VAE as a deep generative model for data-driven protein design blending desirable aspects of variational autoencoders (VAE) and autoregressive (AR) sequence generation. The ProtWave-VAE combines an InfoMax VAE with a dilated convolutional encoder and WaveNet autoregressive decoder and optional semi-supervised regression decoder. This permits model training over unaligned and potentially non-homologous protein families, learning of a meaningful low-dimensional latent space exposing phylogeny and function, reshaping of the latent space and induction of gradients values by semi-supervised training, and autoregressive generation of variable length sequences conditioned on latent vectors and, optionally, N-terminal residues.

We demonstrate and test the predictive and generative capabilities of the ProtWave-VAE model in five applications: (i) learning of biologically meaningful latent space embeddings of four protein families and generative design of novel protein sequences with tertiary structures in close agreement with the natural native folds, (ii) accurate prediction of protein fitness and function in community TAPE and FLIP benchmarks with competitive or superior performance to state-of-the-art architectures, (iii) semi-supervised training over annotated chorismate mutase training data to disentangle functional gradients within the latent space and enable generative design of novel sequences conditioned on high functionality, (iv) C-terminal diversification of synthetic chorismate mutase proteins using N-terminus and latent conditioning, and (v) design and experimental testing of novel SH3 proteins to demonstrate maintenance and elevation of function.

These studies demonstrate the capabilities of the ProtWave-VAE model in data-driven generative protein design. Its capacity for semi-supervised retraining makes it well suited for multi-round protein engineering campaigns within virtuous cycles of model training and synthetic sequence design and testing.^54,55^ Its capacity to learn over unaligned sequence data means that it eschews the need for multiple sequence alignments that can introduce bias into the training data and typically restrict training to homologous protein families. Its capacity for N-terminal conditioning enables directed diversification of the C-terminal region of proteins guided by latent conditioning to introduce or elevate function.

ProtWave-VAE exhibits competitive performance in downstream functional prediction relative to state-of-the-art networks based on large language models, but is much smaller in size, possessing approximately 100-fold fewer trainable parameters. This makes ProtWave-VAE attractive in reducing the cost of training and deployment and accelerating innovation via rapid ablation studies, hyperparameter optimization, and development and testing cycles. The N-terminal conditioning is anticipated to be valuable in protein engineering applications where it is desired to keep part of the protein sequence fixed (e.g., the framework region within an antibody) and generatively design the remainder (e.g., the hypervariable region). Another application is to condition on protein tags (e.g., His-tags or expression-tags) and leverage iterative exploration in the latent space to improve protein expression, stability, or other properties.^8^ A deficiency of the autoregressive nature of ProtWave-VAE is that the conditioning can only be applied unidirectionally, here from N-terminus to C-terminus (or vice-versa). This means it is currently not possible to condition on amino acid residues in arbitrary and non-contiguous regions of the sequence and allow the model to generatively inpaint the remaining residues. A second deficiency is the potential incompatibility of the latent vector and N-terminal conditions in guiding sequence generation. This sets the stage for further innovation to combine latent inference and order-agnostic autoregressive generation for novel protein engineering, and techniques for harmonize multiple conditioning goals. We would also like to apply ProtWave-VAE to the design of larger and more biologically and biomedically relevant proteins, including multi-chain protein complexes with quaternary structure, and also to fields beyond protein engineering that may also benefit from alignment-free, latent-conditioned generative design, including the design of small molecules, nucleic acids, prose, and music.

## Materials and Methods

### Data collection and preparation

Each protein sequence employed in the protein family task, fitness benchmark task, Chorismate mutase unsupervised versus semi-supervised task, and SH3 design task was transformed into one-hot encoded tensors with a length of 21, which includes the 20 amino acid labels and padded tokens. Additional information regarding dataset collection, preprocessing, training protocols, and hyperparameter optimization can be found in the Supporting Information

### Integrating latent-based inference with an autoregressive decoder

To overcome posterior collapse issues and improve variational inference when integrating latent-based inference with autoregressive decoding, we implemented an Information Maximizing VAE model.^24^ Our unsupervised loss function for the ProtWave-VAE model is:

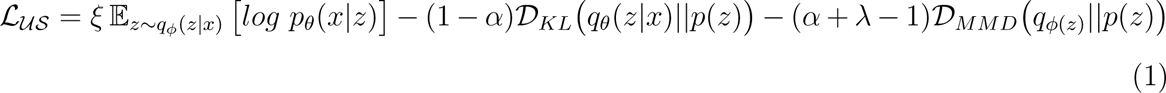

where *p_θ_*(*x|z*) is the decoder model, *D_KL_* is the Kullback-Leibler divergence between the variational posterior approximation *q_φ_*(*z|x*) and normal prior distribution *p*(*z*). The third term *D_MMD_* is the max-mean discrepancy (MMD) that helps penalize the aggregated posterior distribution and improves amortized inference. We introduce an autoregressive decoder emplying a WaveNet-based architecture, where 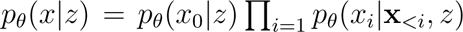. The MMD divergence term becomes 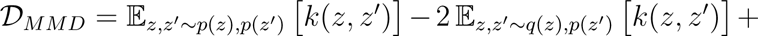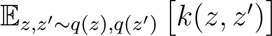, where *k*(*·, ·*) is a positive definite kernel and *D_MMD_* = 0 if and only if *p*(*z*) = *q*(*z*). We choose the Gaussian kernel 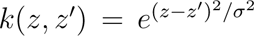 as our characteristic kernel *k*(*·, ·*), and *σ* is a hyperparameter defining the bandwidth of our Gaussian kernel. The prefactor loss weights *ξ*, *α*, and *λ* scale the contribution of the reconstruction loss, weights the mutual information between *x* and *z*, and scales the penalization of MMD divergence.

In Figure 1, the overall architecture of our model is shown, along with three main applications of our approach in protein engineering: alignment-free generation, semi-supervised learning, and C-terminus diversification. The protein sequences, which need not be aligned during either training or deployment, are embedded in a lower-dimensional latent space using a gated dilated convolution neural network encoder *q_φ_*(*z|x*). The decoder (i.e., generator) *p_θ_*(*x|z*) is a WaveNet-based architecture (i.e., gated dilated causal convolution), which samples from the latent space and predicts amino acid residues while conditioning on previous amino acids 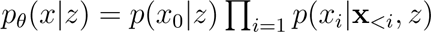. Generally, when using a dilated causal convolution, we use teacher forcing which leverages true previous labeled amino acids as the previous conditional information for predicting the following amino acid label. In contrast to recurrent neural networks or causal masked transformers as the autoregressive decoders, the causal convolutional architectures with teacher forcing allows for fast training with time complexity for the forward pass *O*(1) instead of *O*(*L*), where *L* is the length of the sequence. Recurrent architectures can be prone to vanishing or exploding gradients, whereas this is a deficiency from which convolutional architectures typically do not suffer.

### Extending ProtWave-VAE to a semisupervised learning paradigm

The semi-supervised training objective is the following:

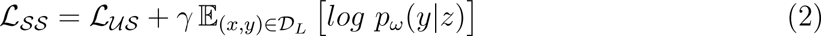

where *p_ω_*(*y|z*) is a regression decoder comprising a simple fully connected neural network parameterized with training parameters *ω*. In practice, we minimize the mean-squared error objective 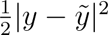, where *y* and 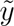 is the ground truth and predicted regression value. The (*x, y*) *∈ D_L_* denotes that the samples which are fed through the supervised branch are only sequences *x* with assay measurements *y*. In the semi-supervised paradigm, the discriminative and generative losses are learned together, allowing reshaping of the latent space for desired engineering tasks and improved control for the design of synthetic sequences with specific functional properties.

### Model architecture, hyperparameter optimization, and training

The encoder architecture included gated nonlinear activations with dilated convolutions and multi-layered perceptrons (MLP) to map the encoder logits to latent vectors. The decoder utilized an autoregressive WaveNet-based architecture with gated activations and dilated causal convolutions. When generating sequences, we first transform the model’s logits into probabilities for each amino acid location, and then select an amino acid label by sampling from the predicted categorical distribution. For certain tasks, such as Chorismate mutase semi-supervised learning and SH3 design, a predictive top model was implemented. This model samples the latent vectors and maps them to protein property predictions using a simple MLP architecture. Hyperparameters were optimized using grid search, and training was conducted using the stochastic gradient descent optimizer Adam.^56^ Full details regarding the architectures, training, and hyperparameter optimization are provided in the Supporting Information and source code can be found here: https://github.com/PraljakReps/ProtWaveVAE.

### ColabFold structure prediction

To predict protein structures for each sequence in the ProtWave-VAE dataset, we employed AlphaFold ColabFold Batch v1.2^31,32^ for AlphaFold2 structure prediction. We generated three structures for each sequence in each task.

### TMalign prediction

We utilized the TMalign algorithm^40^ to calculate the TMscore and heavy-atom root-mean-squared distance (RMSD) between the predictions of natural homolog and design structures. The presented TMscore and RMSD values in the are the mean values of the three ensemble AlphaFold2 ColabFold predictions. The designed sequences with structures that aligned most closely with the natural homologs were considered the best structural matches based on TMscore.

### Sequence novelty

The method for calculating the sequence novelty of the designed samples differs depending on whether the designed sequences are produced by latent-only conditioning or N-terminal and latent conditioning. In the former case, we compute the novelty measurement by determining the minimum Levenshtein distance between the design sample and any natural training sample, then dividing it by the length of the longer sequence in the pair. In the latter case, since we are diversifying a single natural homolog, we calculate the Hamming distance instead of the Levenshtein distance and normalize the Hamming distance by the sequence length of the natural homolog to obtain the sequence dissimilarity. The sequence similarity is commensurately defined as (1 - sequence dissimilarity).

### Gene construction

Experimental protocols follow our previous work. ^14^ *S. cerevisiae* codon-optimized genes coding for all synthetic SH3 proteins were amplified from a mixed pool of oligonucleotide fragments synthesized on microarray chips (Twist). The oligonucleotides corresponding to each gene were designed with primer annealing sites and a padding sequence to make them uniform 250-mer. PCR was performed using KAPA-Hifi polymerase with 1X KAPA HiFi Buffer (Roche), 0.2 mM dNTPs and 1.0 *µ*m of each forward (5’-CCGGTTGTACCTATCGAGTG-3’) and reverse primer (5’-GACCATGCAAGGAGAGGTAC-3’) in 25 *µ*l total volume, with an initial activation (95*^◦^*C, 2 min), followed by 14 cycles of denaturation (95*^◦^*C, 20 s), annealing (65*^◦^*C, 10 s) and primer extension (70*^◦^*C, 10 s). A final extension step (70*^◦^*C, 2 min) was subsequently performed. Amplified products were column purified (Zymo Research), digested with EcoR1 and BamH1, ligated into the digested PRS316 plasmid with N-terminal membrane domain of Sho1,^37^ and transformed into Agilent Electrocompetent XL1-Blues to yield *>*250*×* transformants per gene. The entire transformation was cultured in 50 ml LB media containing 100 *µ*g/ml sodium ampicillin (Amp) at 37*^◦^*C overnight after which plasmids were purified and pooled.

### Sho1 osmosensing high-throughput select-seq assay

Experimental protocols follow our previous work. ^14^ The haploid *S. cerevisiae* strain SS101 was constructed on the W303 background gifted by Wendell Lim (UCSF).^37^ Genetic knock-outs of *Ssk2* and *Ssk22* were created to remove the Sho1-independent branch of the osmoresponse pathway.^57^ The pooled pRS316 plasmids with the SH3 gene library were transformed into SS101 cells using the LiAc-PEG high efficiency transformation protocol.^58^ Plate checks were performed to confirm at least 50 copies of each gene were successfully transformed. Transformed SS101 cells were grown in liquid Sc-Ura media for 24 h (20 mL Sc-Ura media for each 10^8^ total transformed cells) at 30*^◦^*C, and then passaged to 250 mL fresh liquid Sc-Ura media to make OD = 0.05. After another 24 h of growth at 30*^◦^*C, the Sc-Ura culture can be kept at 4*^◦^*C for up to two weeks.

All growth was at 30*^◦^*C on shaker. The stock Sc-Ura culture was transferred to YPD media for a 24 h growth to get the *t*_0_ sample. The culture was diluted every 8 h to keep the cell density below 0.2 OD_600_. A small volume of the *t*_0_ sample was transferred to YPD media supplemented with either (1) no KCl (non-selective) or (2) 1M KCl (selective), and the rest was spun down and mini-prepped to extract plasmids from yeast. Both non-selective and selective cultures were grown for 24 h with OD_600_ maintained under 0.2 to obtain the *t*_24_ samples. The two *t*_24_ samples were span down and minipreped using the same protocol as the *t*_0_ sample.

Plasmids purified from both *t*_0_ and *t*_24_ samples were amplified using two rounds of PCR with Q5 polymerase (New England Biolabs) to add adapters and indices for Illumina sequencing. In the first round the DNA was amplified using primers that add from 6 to 9 random bases (Ns) for initial focusing, as well as part of the i5 or i7 Illumina adapters. Six cycles were used to minimize amplification-induced bias, followed by ampure purification before the second round PCR. In the second round of PCR, the remaining adapter sequence and TruSeq indices were added, where 20 cycles were used. The final products were gel purified (Zymo Research), quantified using Qubit (ThermoFisher), and sequenced in an Illumina MiSeq system with a paired-end 300 cycle kit. Allele counts were obtained using standard procedures. Paired-end reads were joined using FLASH, trimmed to the EcoR1 and BamH1 cloning sites and translated. Only exact matches to the designed genes were counted. Enrichment (*en*) and relative enrichment (*r.e.*) values for each gene *x* of the three growth conditions were defined as,

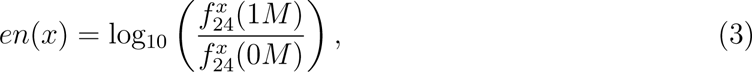

and,

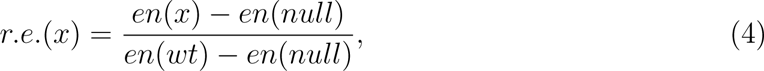

where 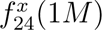 and 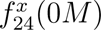 represent the frequency of observing gene *x* after being subjected to a 24 h exposure to 1 M and 0 M, respectively, KCl solution. The wild-type (wt) sequence is the Sho1 gene of *S. cerevisiae* and the null gene is TAGNTAATTTCGGCGTGGGTAT GGTGGCAGGCCCCGTGGCCGGGGGACTGTTGGGCGCCATCTCCTTGCATGCAC CATTCCTTGCGGCGGCGGTGCTCAACGGCCTCAACCTACTACTGGGCTGCTTCCTAATGCAGGAGTCGCATAAGGGAGAGCGTCGAGAT, where the stop codon TAG produces a Sho1 without the C-terminal SH3 domain. A second independent selection assay was performed to ensure reproducibility (Fig. 5A, Fig. S1). The average *en* of the two trials was used to calculate r.e., and the r.e. values of SH3 variants with at least five counts in the 0M at 24 h population in both trials were used for analysis.

## Acknowledgement

We gratefully acknowledge support from the Machine Learning in the Chemical Sciences and Engineering program of The Camille and Henry Dreyfus Foundation (A.L.F.) and grant NIH RO1GM141697 from the National Institutes of General Medical Sciences (R.R.). This work was supported with funding by the University of Chicago Data Science Institute (DSI). This work was completed in part with resources provided by the University of Chicago Research Computing Center. We gratefully acknowledge computing time on the University of Chicago high-performance GPU-based cyberinfrastructure supported by the National Science Foundation under Grant No. DMR-1828629. This material is based upon work supported by the National Science Foundation Graduate Research Fellowship Program under Grant No. 2140001 (N.P.). Any opinions, findings, and conclusions or recommendations expressed in this material are those of the author(s) and do not necessarily reflect the views of the National Science Foundation.

## Abbreviations

AR, autoregressive; CDR, complementary determining region; CM, Chorismate mutase; DGM, deep generative model; DHFR, Dihydrofolate reductase; ELBO evidence lower bound; FLIP, Fitness Landscape Inference for Proteins; MSA, multiple sequence alignment; PCA, principal component analysis; RMSD, root-mean squared distance; SH3, Src homolog 3; TAPE, Task Assessing Protein Embeddings; VAE, variational autoencoder; wt, wild-type.

