## Supplementary Information for "ProtWave-VAE: Integrating autoregressive sampling with latent-based inference for data-driven protein design"

### 1 Data preparation and preprocessing

#### 1.1 Protein family data collection and preprocessing

The datasets used in this study were obtained from Rivoire et al. [11] or from the PFAM database (release 27.0). Accession codes PF00071, PF00186, and PF13354 were used for G proteins, DHFR, and class A $\beta$-lactamases, respectively. For each protein family, a reference sequence/structure was selected, namely rat trypsin (PDB 3TGI) for S1A proteases, human Ras (PDBs 5P21 and 4Q21) for G proteins, E. coli DHFR (PDB 1RX2) for DHFR, and E. coli TEM1 $\beta$-lactamase (PDB 1FQG).

The number of sequences in each dataset was as follows: DHFR (1759 sequences), G-protein (4974 sequences), $\beta$-lactamase (814 sequences), and S1A proteases (1344 sequences). To prepare the sequences for analysis, they were converted into one-hot encoded tensors. In this encoding scheme, each amino acid label corresponds to an index of the one-hot encoded vector along each position of the sequence. The size of the tensor was defined as the maximum length protein homolog in the family. Sequences with less than the maximum sequence length were padded with mask tokens, which have their own one-hot encoded index.

The DHFR, G-protein, beta-lactamase, and S1A proteases datasets were then converted into 2D tensors with sequence lengths of 150, 158, 199, and 205, respectively. The one-hot encoded vector used in the encoding scheme had a length of 21. The datasets can be found here: `https://github.com/PraljakReps/ProtWaveVAE/tree/main/Pfam_analysis/data/protein_families`.

#### 1.2 Fitness prediction benchmarking data collection and preprocessing

The datasets used in this study include the Fitness Landscape Inference for Proteins (FLIP) and the Tasks Assessing Protein Embedding (TAPE). The FLIP datasets consist of two benchmarking tasks (AAV and GB1) and were obtained from FLIP benchmarks [4]. The TAPE datasets also include two benchmarking tasks and were obtained from TAPE benchmarks [9].

The AAV benchmarking task involves a mutational screening landscape of VP-1 AAV proteins [2, 19] (UniProt Accession P03135). Sequences were mutagenized along a 28-amino acid window from position 561 to 588 of VP-1, and the resulting variants were tested for fitness, with between 1 and 39 mutations. The VP-1 AAV benchmark task has seven dataset splits: (1) sampled, (2) sampled-designed, (3) design-sampled, (4) train on single mutants, (5) train on single and double mutants, (6) train on mutants with up to seven changes, and (7) train on low fitness, test on high.

The GB1 benchmarking task involves an exhaustive, combinatorial, and highly epistatic mutational landscape, ranging from variants with fewer mutations to predict activity of variants with more mutations [18]. The number of fitness measurements was 149,361 out of 160,000 possible combinations of mutations at four positions. The GB1 benchmark task also has seven dataset splits: (1) sampled, (2) train on single mutants, (3) train on single and double mutants, (4) train on single, double, and triple mutants, and (5) train on low fitness, test on high.

The TAPE benchmarking tasks include a green fluorescence protein (GFP) and stability measurements of candidate proteins. For the fluorescence regression predictions, a Deep Mutational Scanning (DMS) approach was used to characterize the local genotype-to-phenotype mapping of a single protein [14]. The training and validation sets include mutants with only Hamming distance 3 from the original protein, while the testing set includes mutants with Hamming distances 4-15. In contrast, the stability task [12] has a training and validation set containing proteins from four rounds of experimental data measuring candidate proteins, while the testing set consists of mutant variants of seventeen 1-Hamming distance neighborhoods. The FLIP datasets can be found here: https://benchmark.protein.properties/, while the TAPE datasets can be found here: https://github.com/songlab-cal/tape#data.

##### 1.3 Chorismate mutase data collection and preprocessing

The dataset used in this study was obtained from Russ et al. [13]. For each protein family, four CM atomic structures (PDB entries 1ECM, 2D8E, 3NVT, 1YBZ) were selected as reference sequences/structures to build the multiple sequence alignment. The 1ECM PDB corresponds to the wild-type Chorismate mutase *E. coli*. The training dataset consists of natural homolog sequences, with 1130 sequences, while the testing dataset consists of synthetic natural homolog sequences, with 1618 sequences. Each protein sequence has a corresponding fitness value quantified by the normalized relative enrichment (r.e.).

To prepare the sequences for analysis, they were converted into one-hot encoded tensors, with each amino acid label corresponding to an index of the one-hot encoded vector along each position of the sequence. The size of the tensor was defined as the maximum length protein homolog in the family. Sequences with less than the maximum sequence length were padded with mask tokens, which have their own one-hot encoded index.

The CM dataset was then converted into a 2D tensor with a sequence length of 96. The one-hot encoded vector used in the encoding scheme had a length of 21. The dataset can be found here: `https://github.com/PraljakReps/ProtWaveVAE/tree/main/Pfam_analysis/data/protein_families`.

##### 1.4 SH3 protein data collection and preprocessing

The datasets used in this study were obtained from [7], and consist of SRC homolog 3 (SH3) homologs. The dataset contains various natural paralogs, Sho1 orthologs, and synthetic domains with fitness measurement scores, quantified by the relative enrichment (r.e.). The dataset size is 17,218 sequences, which includes both natural and synthetic sequences.

To prepare the sequences for analysis, the input data tensor was set to a length of 82, corresponding to the maximum sequence length within the dataset. Shorter sequences were padded with mask tokens at the ends. Each sequence was then converted into 2D tensors using a one-hot encoded transformation, such that each amino acid and masked padded token corresponded to a one-hot encoded index.

The number of sequences with fitness measurements is 14,768. During stratified cross-validation, we set k = 5 and split the dataset into five partition bins, such that the training and validation set sizes are 80% and 20%, respectively. The dataset can be found here: `https://github.com/PraljakReps/ProtWaveVAE/tree/main/SH3_design_project/data`.

#### 2 Model architecture and hyperparameterization

##### 2.1 Gated dilated encoder

The primary hyperparameters for the gated dilated convolutional encoder network $q_\phi(z|x)$ include encoder depth, input channel depth for the initial convolutional layer, hidden channel depth for the subsequent convolutional layers, kernel size for each convolutional layer, and the number of fully connected linear layers before transitioning the encoder outputs into the latent space. We increased the dilation window for the convolutional layer by $2^i$, based on the encoder layer depth (where $i$ is the depth index of the encoder), creating what is known as dilated convolution layers.

For each protein family task, the encoder depth was chosen to ensure that the original input sequence remained greater than length 0 after the convolutional layers reduced its length. Consequently, hyperparameter optimization for the encoder depth was not necessary, as it depended on the input sequence length `compute_max_enc_depth`. Following the approach used in WaveNet [16] and PixelCNN [17], we employed two sets of dilated convolutions: one for signal activations and another for gated activations.

Our nonlinear activation function was a linear gated activation function [5], which multiplied the outputs of the signal and gated convolution layers after applying a sigmoid function onto the gated convolution outputs. This enabled the encoder to incorporate a memory gate similar to LSTMs and GRUs [6, 3]. A batch normalization layer was applied after each gated activation operation. Once the dilated convolution layers processed the hidden representations, signal and gate convolution layers with kernel size 1 and channel depth 1 were applied to the outputs. Subsequently, a gated activation operation and a batch normalization layer were added.

Next, we applied $k$ fully connected layers, using leaky ReLU activation functions with an $\alpha$ hyperparameter of 0.1. The final outputs were then passed through a variational mean linear layer and a variational variance module, which comprised a linear layer followed by a softplus activation function. Ultimately, the final variational mean and variational variance outputs were combined using the reparameterization trick (reparam_trick). This involved sampling Gaussian noise from a normal distribution $\epsilon$, multiplying it with the square root of the variational variance, and adding the variational mean to produce the latent vector. The forward pass for the encoder network is shown in pseudocode (Algorithm 1). The source code and class object of the encoder network is in PyTorch and can be found in the following link: https://github.com/PraljakReps/ProtWaveVAE/blob/main/SH3_design_project/source/model_components.py.

```
def compute_L_out(L_in, dilation, kernel_size=3, padding=0, stride=1):
    return (L_in + 2*padding - dilation * (kernel_size - 1) - 1) / stride + 1

def compute_max_enc_depth(max_protein_length):

    # loop over all possible hyperparameter options
    for max_encoder_dilation in range(1, 100):
        dilation = [2**ii for ii in range(max_encoder_dilation)]
        L = max_protein_length
        """
        check whether the output length after dilation convolution
        layer is greater than 0.
        """
        try:
            for i in dilations:
                # output sequence length after conv. operation
                L_out = compute_L_out(L_in=L, dilation=ii)
                # make sure output sequence is longer than 0
                if L_out <= 0:
                    raise StopIteration
                else:
                    pass

        except StopIteration:
            # return maximum allowed dilation value
            return max_encoder_dilation - 1

def reparam_trick(z_mean, z_var):
    # compute the standard deviation from the variance
    std = z_var.sqrt()
    # sample from normal distribution with pytorch programming
    eps = torch.rand_like(std)
    return z_mean + std * eps
```

**Algorithm 1** Gated dilated convolution encoder network $q_\phi(z|x)$

```
Input: {
x, encoder_depth, num_fc, sigmoid, lrelu, initial_conv_layer,
signal_dilated_conv_layers, gate_dilated_conv_layers,
batc_norm_layers, final_signal_conv_layers, final_gate_conv_layers,
fc_layers, mean_z_layers, var_z_layers
}
// *Description of inputs.*
// x --> input sequence data
// encoder_depth --> number of dilated convolution layers
// num_fc --> number of fully connected linear layers
// sigmoid --> sigmoidal activation function
// lrelu --> leaky ReLU activation function
// initial_conv_layer --> first convolution layer with kernel size 1
// signal_dilated_conv_layers --> list of layers along the signal network path,
   consisting of dilated convolutions with increasing dilations by factor of 2
// gate_dilated_conv_layers --> list of layers along the gated network path,
   consisting of dilated convolutions with increasing dilations by factor of 2
// batch_norm_layers --> list of batch normalization layers
// batch_norm_layers --> list of batch normalization layers
// final_signal_conv_layers --> single convolution layer with kernel size 1 and depth
   channel 1
// final_gate_conv_layers --> single convolution layer with kernel size 1 and depth
   channel 1
// final_gate_conv_layers --> single convolution layer with kernel size 1 and depth
   channel 1
// fc_layers --> fully connected linear layers
// mean_z_layers --> fully connected linear layers mapping encoder outputs to latent
   mean
// var_z_layers --> fully connected linear layers mapping encoder outputs to latent
   variance
batch_size, input_channels, max_protein_length = x.shape
encoder_depth = compute_max_enc_depth(
   max_protein_length=max_protein_length
   ) // Determine encoder depth based on protein sequence length
h = init_conv_layer(x)// 1x1 convolution operation
h = batch_norm[i](h)
for i ← 1 to encoder_depth
  // signal and dilated convolution layer operation
  h_signal=signal_dilated_conv_layers(dilation=2^(i−1))[i](h)
  h_gate=gate_dilated_conv_layers(dilation=2^(i−1))[ii](h)
  h = h_signal * sigmoid( h_gate)
  h = batch_norm_layers[ii+1](h)
  end for
// final single no-dilated convolution layer to map output depth channels to 1
h_signal = final_signal_conv_layer(h)
h_gate = final_gate_conv_layer(h)
// apply gated activation function
h_out = h_signal * sigmoid(h_gate)
h_out = final_gate_conv_layer(h_out)
for i ← 1 to num_fc
  h_out = fc_layers[ii](h_out)
  h_out = lrelu(h_out)
  end for
z_mean, z_var = mean_z_layer(h_out), var_z_layer(h_out)
// infer latent variables using reparameterization trick
z = reparam_trick(z_mean, z_var) Output: z, z_mean, z_var
```

#### 2.2 WaveNet decoder with latent conditioning

The WaveNet architecture is based on the work of van Oord et al. [16], utilizing dilated causal convolution layers. Our approach differs in that we employ latent variables for conditional inference rather than one-hot encoded class variables. In accordance with the class conditional WaveNet, we upsample our latent variables through a linear layer, enabling us to extend the latent vector to the sequence data dimension's length. This is referred to global conditioning [16]. No activation function is applied during this linear upsampling for the latent variable, and the latent linear upscaling operation for global conditioning is the following:

$$Z = V_k^T \ z \tag{1}$$

where $V_k$ is learnable linear project, $z$ is the inferred latent low-dimensional varible, $Z$ is the vector broadcast over the sequence position dimension. This operation is implemented by `latent_cond_net`() nn.module, and the source code is https://github.com/PraljakReps/ProtWaveVAE/blob/01cecd054be7ca77540b23d19aeea5a3b9012d79/SH3_design_project/source/wavenet_decoder.py#L23.

The WaveNet decoder has two primary components: the main module, which is composed of dilated causal convolutions, gated activation functions, residual connections, and skip connections; and the top head, which is inspired by van Oord et al.'s architecture [16] and features two convolution layers (non-dilated) with kernel size 1 and a ReLU activation function. The second component generates logits that is converted to the amino acid class distribution using a softmax function. The main hyperparameter optimization for the first component involves input channel depth, output channel depth, and the number of dilation rates (i.e., the WaveNet's depth). The causal dilation convolutions' long-range reach is determined by the last hyperparameter. The top model's hyperparameters include input channel depth, output channel depth, and hidden state depth. The output channel depth is set to the WaveNet's input channel depth, which corresponds to the one-hot encoded vector's length (20 amino acids plus 1 padded token).

The WaveNet decoder processes the upsampled latent variable and input sequence. The first layer applies a convolution layer with kernel size 1 to the input sequence, creating an amino acid embedding layer $h = W_{k=1} \ x$, where $W_{k=1}$ is the 1×1 kernel with learnable weights. Subsequently, the causal dilated convolution is employed in two separate modules for signal and gated representations. At the same layer depth as the dilated convolutions, a vanilla convolution layer with kernel size 1 is also applied to the upsampled latent variable, mapping input depth 1 to output depth as per the hyperparameter $C_{out}$. The hidden representation outputs from the signal causal dilated convolution are then summed with the outputs from the signal latent convolution, and the outputs from the gated causal convolution are summed with those from the latent gated convolutions. A sigmoid function is applied as a gating function to the gated output representations, and these values are multiplied with the signal hidden representations. After completing this process for a given dilated causal convolution layer, a convolution layer corresponding to the skip module and another convolution layer for the residual module, both with kernel size 1, are applied. Thus, in terms of mathematical expression, the layer operations are the following:

$$h = \big(W_{a,k} * h + \tilde{W}_{a,k=1} * Z\big) \odot \sigma\big(W_{g,k} * h + \tilde{W}_{g,k=1} * Z\big) \tag{2}$$

where first term in the parentheses corresponds to linear activations $a$ while the second term in the parentheses corresponds to the gated inputs to the sigmoidal function $\sigma$, acting as a gated function and outputting gated activations $g$. The learnable kernel weight for the sequence representations and latent variable representations are $W_{*,k}$ and $\tilde{W}_{*,k=1}$. The variables $h$ and $Z$ corresponds to the hidden representations of the input protein sequence and upsampled global latent conditioning.

The skip output representations are accumulated (i.e., summed) after each dilated causal convolution layer, are implemented with a linear convolution with kernel size 1, and independently applied relative the to residual layer; and thus the mathematical operation for this layer are the following:

$$h^l_{skip} = V^{skip}_{l,k=1} * h \tag{3}$$

$$h_{cumm-skip} \mathrel{+}= h^l_{skip} \tag{4}$$

where $h_{cumm-skip}$ is the cumulative skip representations of all the skip output representations from the $l$ dilated causal convolution layers applied to the input sequence. The output representations from the residual module's convolution layers are added to the embedded input sequence, which is then fed back into the next dilated causal convolution layer with an increased dilation scale by $2^i$, where $i$ is the depth index of the dilated causal convolution layer. The mathematical operations are then the following:

$$h_{res} = V^{res}_{l,k=1} * h \tag{5}$$

$$h = h + h_{res} \tag{6}$$

The top model exclusively utilizes the accumulated skip connections as input and processes these hidden representations using a ReLU function, a convolution layer with kernel size 1, a second ReLU activation function, and finally a second convolution layer with kernel size 1. The top model generates logits that can be converted into probabilities using a softmax function. Thus, the final sequence probabilities using the top model is following:

$$\begin{aligned} h^1_{top-model} &= Conv\big(ReLU(h_{cumm-skip})\big) \\ h^2_{top-model} &= Conv\big(ReLU(h^1_{top-model})\big) \end{aligned} \tag{7}$$

$$p(x|z) = softmax(h^2_{top-model}) \tag{8}$$

where $ReLU$ is elementwise nonlinear activation function and $Conv$ is a linear learnable affine transformation with bias and kernel size 1. The softmax layer converts the network logits into probabilties using the following function $softmax(x_i) = \frac{exp(x_i)}{\sum_j exp(x_j)}$

The source code for the class objects, written in PyTorch, can be found via this https://github.com/PraljakReps/ProtWaveVAE/blob/01cecd054be7ca77540b23d19aeea5a3b9012d79/SH3_design_project/source/wavenet_decoder.pyL23, with the pseudocode provided below. The pseudocode for the upsampling latent network in PyTorch, comprising a solitary linear layer, is displayed below.

```
class latent_cond_net(nn.module):

    def __init__(
            self,
            z_dim: int,
            output_shape: any=(1,max_protein_length)
    ):

    self.input_shape = z_dim
    self.output_shape= output_shape
    self.linear = nn.Linear(
                    self.input_shape,
                    np.prod(self.output_shape),
                    bias=False
    )

    def forward(self, z: torch.FloatTensor) -> torch.FloatTensor:
        return self.linear(z).view(-1,1,self.output_shape[-1])
```

The subsequent PyTorch pseudocode represents the WaveNet decoder, encompassing numerous causal dilation convolutions, latent convolution layers, a skip module with convolution layers, and a residual module with convolution layers (see https://github.com/PraljakReps/ProtWaveVAE/blob/01cecd054be7ca77540b23d19aeea5a3b9012d79/SH3_design_project/source/wavenet_decoder.py#L98).

```
class WaveNet_head(nn.module):

    def __init__(
            self,
            device: str,
            C_in: int,
            C_out: int,
            num_dilation_rates: int
            **kwargs
            ):

        """
        initialize layers and variables ... see source code for more details:
        https://github.com/PraljakReps/ProtWaveVAE/blob/main
        /SH3_design_project/source/wavenet_decoder.py
        """

    def forward(
        self,
        x: torch.FloatTensor,
        z: torch.FloatTensor
    ) -> (
            torch.FloatTensor,
            torch.FloatTensor
    ):

    h = self.emb_conv(x) # 1x1 initial conv operation

    # create residual and skip variables
    orig_x = h.clone()
    h_cum_skip = 0

    # loop over the number of dilation rates (increase the mode's receptive field)

    for ii in range(self.num_rates):

        # conditional operation for signal path: W_s*x + \tilde{W}_s*x
        h_signal = self.signal_conv[ii](h) + self.latent_signal_conv[ii](z)

        # conditional operation for gate path: W_g*x + \tilde{W}_g*x
        h_gate = self.gate_conv[ii](h) + self.gate_signal_conv[ii](z)

        # gate operation
        h = h_signal * self.sigmoid(h_gate)

        # output skip and residual representations
```

```
        h_skip = self.skip_blocks[ii](h)
        h_res = self.residual_blocks[ii](h_)

        # residual path
        if ii == 0:
            h = orig_x + res
        else:
            h = res + h

        # cummulate all skip connection representations
        h_cum_skip = h_cum_skip + h_skip

    return (
            h,
            h_cum_skip
    )
```

The final PyTorch pseudocode pertains to the TopHead of the WaveNet decoder, which produces the ultimate logits before utilizing a softmax function to transform them into amino acid class probabilities. This component features two straightforward convolution layers and two ReLU functions, while solely accepting cumulative skip outputs as input (see https://github.com/PraljakReps/ProtWaveVAE/blob/01cecd054be7ca77540b23d19aeea5a3b9012d79/SH3_design_project/source/wavenet_decoder.py#L222).

```
class Top_head(nn.module):

    def __init__(
            self,
            C_in: int,
            C_out: int,
            hidden_state: int
        ):

        self.conv1 = nn.Conv1d(C_in, hidden_state, 1, bias=True)
        self.conv2 = nn.Conv1d(hidden_state, hidden_state, 1, bias=True)

        self.relu = torch.nn.ReLU()

    def forward(
            self,
            h_cum_skip: torch.FloatTensor
        ) -> torch.FloatTensor:
        """
        same architecture as the diagram in Figure 1 (main text)
        and Figure 4 in van der Oord et al. arXiv 2016.
        """

        h = self.conv1(self.relu(h_cum_skip))
        logits =  self.conv2(self.relu(h))

        # output only logits and not yet probs.
```

```
        return logits
```

Forward pass of the latent conditional WaveNet deocder in psuedocode (see https://github.com/PraljakReps/ProtWaveVAE/blob/01cecd054be7ca77540b23d19aeea5a3b9012d79/SH3_design_project/source/wavenet_decoder.py#L303).

**Algorithm 2** Forward pass for latent conditioned WaveNet decoder $p_\theta(x|z)$

**Input:** x, z, softmax, latent_cond_net, WaveNet_head, Top_head

```
// *Description of inputs.*
// x --> input sequence data
// z --> inferred latent variable data
// latent_cond_net --> upsampling linear layer for the latent variables
// WaveNet_head --> WaveNet causal dilated convolution component
// Top_head --> Top model component, consisting of convolution layers and outputting
   amino acid logits
z_upsampled = latent_cond_net(z)
tot_cum_skip = 0 // Initialize the cumulative skip representations
h, h_cum_skip = WaveNet_head(x, z_upsampled)
tot_cum_skip += h_cum_skip // Cumulate WaveNet's skip representations
logits = Top_head(tot_cum_skip) // final operation to compute amino acid logits
p(x|z) = softmax(logits)
```

**Output:** p(x|z)

#### 2.3 Discriminate top model over the latent space for semi-supervision

In our semi-supervised learning implementation, we utilize a top model that samples from the latent variables and predicts fitness regression values $y$ and/or classifies sequences as functional or nonfunctional. For regression tasks, we employ the mean-squared error as our loss objective. Conversely, for classification tasks, we use the binary cross-entropy loss, $l(y, \tilde{y})$, where the loss value for a given sample sequence $n$ is defined as $l_n = y_n * \log \tilde{y}_n + (1 - y_n) * \log(1 - \tilde{y}_n)$. Here, $\tilde{y}_n$ represents the classification prediction probability, while $y_n$ denotes the ground truth label (0 or 1).

The model architecture remains consistent, regardless of whether the task is regression or classification. It consists of hidden modules, with each module containing the following layers:

$$h = Dropout(SiLU(LayerNorm(Linear(z)))) \tag{9}$$

Dropout is a widely used regularization layer for neural networks [15]. SiLU represents a nonlinearity function, LayerNorm is a layer normalization layer [1], and Linear refers to a simple fully connected linear layer. The primary hyperparameters include `num_layers`, which defines the number of hidden modules to stack, and `hidden_width`, which sets the hidden width size of the linear layers. The hidden output representations $h$ are then fed into a final linear layer for either regression or classification predictions. The classification path concludes with a sigmoidal function that maps the values to a binary probability, while the regression path does not require any final nonlinear layer since the outputs are continuous values. The hidden module's architecture is denoted as `TopModel_layer`, while the overall neural network architecture $p_\omega(y|z)$ is illustrated in object `Decoder_re`. The model and hidden module's source code can be found at this https://github.com/PraljakReps/ProtWaveVAE/blob/01cecd054be7ca77540b23d19aeea5a3b9012d79/SH3_design_project/source/model_components.py.

The hidden blocks found along the top model discriminator $p_\omega(y|z)$, which consist of linear layer followed by a layer normalization layer [1], then followed by a SiLU nonlinearity and dropout regularization layer [15]. The subsequent pytorch psuedocode represents these hidden block modules called `TopModule_layer`.

The top model discriminator's hidden blocks, $p_\omega(y|z)$, consist of a linear layer followed by a layer normalization layer [1], a SiLU nonlinearity, and a dropout regulation layer [15]. The subsequent PyTorch pseudocode represents these hidden block modules, referred to as TopModule_layer.

```
class TopModel_layer(nn.module):

    def __init__(
            self,
            in_width: int,
            out_width: int,
            p: float=0.1
    ):

    self.in_width = in_width
    self.out_width = out_width
    self.p = p

    self.hidden_layer = nn.Sequential(
                nn.Linear(self.in_width, self.out_width),
                nn.LayerNorm(self.out_width),
                nn.SiLU(),
                nn.Dropout(p=self.p)
    )

    def forward(self, x: torch.FloatTensor) -> torch.FloatTensor:
        return self.hidden_layer(z)
```

We present the entire neural model for the top discriminate model $p_\omega(y|z)$. This architecture encompasses both regression and classification tasks. However, the network can be easily adapted to cater to individual discriminate tasks. The subsequent PyTorch pseudocode represents these hidden block modules, denoted as Decoder_re. The source code can be found https://github.com/PraljakReps/ProtWaveVAE/blob/01cecd054be7ca77540b23d19aeea5a3b9012d79/SH3_design_project/source/model_components.py#L279.

```
class Decoder_re(nn.module):

    def __init__(
            self,
            num_layers: int,
            hidden_width: int,
            z_dim: int,
            num_classes: int,
            p: float=0.1
    ):

    # general variables used throughout the model
    self.num_layers = num_layers
    self.hidden_width = hidden_width
    self.z_dim = z_dim
    self.num_classes = num_classes
    self.p = p
```

```python
"""
Optional: Here, we will present a discriminative top model
that has a regression and classification discriminator. However,
this component can easily be modified for either regression or
classification such that the neural network models p(y|z).
"""
self.reg_model_layers = nn.ModuleList()
self.class_model_layers = nn.ModuleList()

for ii in range(num_layers):

    if ii == 0:
        # regression path
        self.reg_model_layers.append(
                TopModel_layer(
                                in_width=self.z_dim,
                                out_width=self.hidden_width,
                                p=self.p
                )
        )

        # classification path
        self.class_model_layers.append(
                TopModel_layer(
                                in_width=self.z_dim,
                                out_width=self.hidden_width,
                                p=self.p
                )
        )

    else:
        # regression path
        self.reg_model_layers.append(
                TopModel_layer(
                                in_width=self.hidden_width,
                                out_width=self.hidden_width,
                                p=self.p
                )
        )

        # classification path
        self.class_model_layers.append(
                TopModel_layer(
                                in_width=self.hidden_width,
                                out_width=self.hidden_width,
                                p=self.p
                )
        )
    # final output layers
    self.output_class_layer = nn.Linear(self.hidden_width, self.num_classes)
    self.output_reg_layer = nn.Linear(self.hidden_width, self.num_classes)
```

```python
    def reg_forward(self, z: torch.FloatTensor) -> torch.FloatTensor:
        for layer in self.reg_model_layers:
            z = layer(z)
        return self.output_reg_layer(z)

    def class_forward(self, z: torch.FloatTensor) -> torch.FloatTensor:

        for layer in self.class_model_layers:
            z = layer(z)
        return self.sigmoid(self.output_class_layer(z))

    def forward(self, z: torch.FloatTensor) -> (
            torch.FloatTensor,
            torch.FloatTensor
        ):
        y_reg = self.reg_forward(z)
        y_class = self.class_forward(z)
        return (
                y_reg,
                y_class
        )
```

We demonstrate the forward pass of the latent discriminative top model $p_\omega(y|z)$ in pseudocode (refer to the https://github.com/PraljakReps/ProtWaveVAE/blob/01cecd054be7ca).

**Algorithm 3** Forward pass for latent discriminative top model $p_\omega(y|z)$

**Input:** z, Decoder_re

```
// *Description of inputs.*
// z --> inferred latent variable data
// Decoder_re --> discriminative regression and classification model
y_pred_R, y_pred_C = Decoder_re(z) // model predictions p(y|z)
```

**Output:** y_pred_R, y_pred_C

#### 2.4 Model architecture and hyperparameter optimization for protein family task

The task of the protein family is to evaluate the ability of ProtWave-VAE to generate biologically meaningful latent representations for a given protein family, without any supervision (i.e., unsupervised). Additionally, since the model is generative, this task tests its ability to generate novel sequences that are indistinguishable from the protein family in terms of tertiary structure, using known PDB structure and ColabFold structure predictions. Therefore, the architecture of ProtWaveVAE for protein family tasks (specifically Gprotein, DHFR, S1A, and lactamase families) consists of the gated dilated encoder $q_\phi(z|x)$ with the latent conditioned WaveNet decoder $p_\theta(x|z)$, since it is solely unsupervised. The loss objective is the unsupervised loss, which includes the negative log-likelihood, Kullback Leibler divergence between the posterior and prior latent distribution, and the max-mean discrepancy between the aggregated posterior and prior latent distribution. Since each family has different maximum sequence lengths, the encoder depth is determined based on the maximum possible depth, or the number of dilated convolutions that can be applied onto the input sequence before compressing the sequence length below 0. The hyperparameters that were optimized are the latent space dimension `z`, channel depth of dilated convolution in the WaveNet decoder `whs`, channel depth of the 1x1 convolution layers for WaveNet TopHead `hhs`, channel depth of the encoder convolution layers `C_out`, number of fully connected layers along the encoder path `num_fc`, number of dilated causal convolutions along the decoder path `ndr`, prefactor weight for the KL divergence loss term `KL_weight`, prefactor weight for the negative log-likelihood term `NLL_weight`, and the prefactor weight for the max-mean discrepancy term `MMD_weight`. These hyperparameters were optimized in order, and the optimal values were determined by the optimal negative log-likelihood loss while maintaining excellent max-mean discrepancy regularization loss on a random hold-out that consisted of 20% of the original protein family dataset. The optimization was performed over the domains `z`$\in [1, 20]$, `whs`$\in \{32, 64, 128, 256, 512\}$, `hhs`$\in \{32, 64, 128, 256, 512\}$, `C_out`$\in \{32, 64, 128, 256, 512\}$, `num_fc`$\in$[0,4], `ndr`$\in$[1,10], `KL_weight`$\in \{0.7, 0.75, 0.8, 0.85, 0.9, 0.95, 0.99\}$, `NLL_weight`$\in \{0.1, 0.5, 1.0, 5.0, 10.0, 50.0, 100.0\}$, and `MMD_weight`$\in \{1.0, 2.0, 5.0, 10.0, 20.0, 50.0, 100.0\}$. The training consisted of using an Adam optimizer with a learning rate equal to $1 \times 10^{-4}$. The minibatch size was 256, and the number of epochs was 500. The model was trained on unaligned sequences for each protein family.

The final optimized hyperparameter configurations for the G-protein, DHFR, S1A, and lactamase families are presented below. For G-protein, the model configuration is `z`=6, `whs`=128, `hhs`=512, `C_out`=32, `num_fc`=3, `ndr`=5, `KL_weight`=0.99, `NLL_weight`=1.0, `MMD_weight`=5. For DHFR, the model configuration is `z`=6, `whs`=128, `hhs`=256, `C_out`=32, `num_fc`=3, `ndr`=5, `KL_weight`=0.99, NLL_ weight=1.0, `MMD_weight`=10. For S1A, the model configuration is `z`=4, `whs`=128, `hhs`=512, `C_out`=256, `num_fc`=3, `ndr`=5, $\alpha$=0.99, $\xi$=10.0, $\lambda$=1. For lactamase, the model configuration is `z`=3, `whs`=128, `hhs`=512, `C_out`=512, `num_fc`=1, `ndr`=7, `KL_weight`=0.99, `NLL_weight`=100.0, `MMD_weight`=10. These models were k-fold cross-validated with k=5 and found that the average negative log-likelihood and max-mean discrepancy were consistent across different folds and close to the training values (result spreadsheets found https://github.com/PraljakReps/ProtWaveVAE/tree/01cecd054be7ca77540b23d19aeea5a3b9012d79/Pfam_analysis/outputs/train_sess/pfam), indicating that the models are not overfitting on the training set. Next, after verifying that the model is not overfitting on the given random training set split, the final model was trained on the entire dataset using the same number of epochs (500). The work presented in protein inference and generative design here utilized these models trained on the whole protein family dataset.

#### 2.5 Model architecture and hyperparameter optimization for benchmark fitness task

The fitness and function benchmarking tasks aim to assess the ability of ProtWave-VAE to predict regression values for various extrapolative problems. By leveraging semi-supervision, we introduce a second decoder head that incorporates a regression model, allowing us to sample latent variables $z$ and predict functional/fitness continuous values $y$. This task tests the model's capacity to use the inferred latent variables for more than simply generative design, but for extrapolative regression prediction, and more importantly, compare it against state-of-the-art models, including large language models (e.g., ESM-1 [10]). The architecture of ProtWave-VAE for the fitness benchmarking tasks consists of a gated dilated encoder $q_\phi(z|x)$, a latent-conditioned WaveNet decoder $p_\theta(x|z)$, and a discriminative multi-layer perceptron model $p_\omega(y|z)$. The decoder component that is implemented for regression follows the architecture `Decoder_re`.

The following hyperparameters were optimized for the ProtWave-VAE model: (1) latent space embedding size `z_dim`, (2) number of kernel channels for the dilate convolutions along the encoder path `C_out`, (3) number of fully connected layers at the end of the encoder path `num_fc`, (4) number of channels implemented in the dilated causal convolutions along the WaveNet decoder path `whs`, (5) number of channels implemented in the 1x1 convolutions along the WaveNet TopHead decoder path `hhs`, (6) number of dilated causal convolutions used `ndr`, (7) number of discriminative hidden module layers `disc_num_layers`, (8) width of the hidden linear layers along the discriminative decoder `hidden_width`, (9) dropout probability for the discriminative decoder path `p`, (10) negative log-likelihood prefactor weight `NLL_weight`, (11) Kullback-Leibler divergence prefactor weight `KL_weight`, (12) max-mean discrepancy prefactor weight `MMD_weight`, and (13) discriminative likelihood prefactor weight gamma_weight. These hyperparameters were optimized for each of the following four protein families: AAV-capsid task (FLIP), GB1 task (FLIP), GFP task (TAPE), and stability task (TAPE). The hyperparameters were optimized sequentially, with the optimal values chosen based on the optimal spearman $\rho$ correlation and mean square error loss while maintaining good negative log-likelihood loss and excellent max-mean discrepancy regularization loss. For FLIP benchmarks, we hyperparameter optimized over the random split train/valid split and used those hyperparameters for the remaining benchmark splits. While for for the TAPE benchmarks, we simply hyperparameter optimized based on the chosen validation set by the TAPE authors.

The optimization process covered the following domains: (1) z_dim$\in [1, 20]$, (2) C_out$\in \{32, 64, 128, 256, 512\}$, (3) whs$\in \{32, 64, 128, 256, 512\}$, (4) hhs$\in \{32, 64, 128, 256, 512\}$, (5) num_fc$\in [0, 5]$, (6) ndr$\in \{2, 4, 6, 8, 10\}$, (7) disc_num_layers$\in 1, 2, 3, 4, 5$, (8) hidden_width$\in \{2, 5, 10, 20, 50, 100, 200, 500, 1000\}$, (9) p$\in \{0.0, 0.1, 0.2, 0.3, 0.4, 0.5\}$, (10) NLL_weight$\in \{0.1, 0.5, 1.0, 5.0, 10.0, 50.0, 100.0\}$, (11) KL_weight$\in \{0.8, 0.85, 0.9, 0.95, 0.99\}$, (12) MMD_weight$\in \{1.0, 2.0, 5.0, 10.0, 20.0, 50.0, 100.0, 200.0, 500.0, 1000.0\}$, and (13) gamma_weight $\in \{0.1, 0.5, 1.0, 5.0, 10.0, 50.0, 100.0\}$. An Adam optimizer with a learning rate of $1 \times 10^{-4}$ was used during training, and unaligned sequences were employed for all benchmarking tasks. For the TAPE benchmark tasks, the training set was split into a train/validation set, and the previously mentioned metrics were optimized on the validation set. The AAV capsid task used an epoch of 500 and a batch size of 256, while the GB1 task used an epoch of 500 and a batch size of 512. For the GFP task, a batch size of 256 and an epoch of 300 were used, while for the stability task, a batch size of 256 and an epoch of 300 were used.

The final optimized hyperparameter configurations for the AAV capsid, GB1, GFP, and stability tasks are presented below. For AAV capsid, the model configuration is z_dim = 6, C_out = 128, whs = 128, hhs = 256, num_fc = 3, ndr = 8, disc_num_layers = 1, hidden_layers = 50, p = 0.3, NLL_weight = 100.0, KL_weight = 0.99, MMD_weight = 10.0, and gamma_weight = 10. For GB1, the model configuration is z_dim = 3, C_out = 128, whs = 256, hhs = 512, num_fc = 3, ndr = 6, disc_num_layers = 5, hidden_layers = 20, p = 0.0, NLL_weight = 50.0, KL_weight = 0.99, MMD_weight = 10.0, and gamma_weight = 10. For GFP, the model configuration is z_dim = 4, C_out = 512, whs = 256, hhs = 256, num_fc = 2, ndr = 8, disc_num_layers = 2, hidden_layers = 500, p = 0.3, NLL_weight = 10.0, KL_weight = 0.99, MMD_weight = 20.0, and gamma_weight = 50. For stability, the model configuration is z_dim = 11, C_out = 512, whs = 64, hhs = 128, num_fc = 0, ndr = 10, disc_num_layers = 4, hidden_layers = 50, p = 0.3, NLL_weight = 0.50, KL_weight = 0.99, MMD_weight = 100.0, and gamma_weight = 5.0, while training configuration was epochs equal to 2000 and batch size equal to 512. After finding the optimal hyperparameter configuration for which task, we used these hypeparameters on the train/test splits defined for the benchmarking tasks.

#### 2.6 Model architecture and hyperparameter optimization for Chorismate Mutase task

To assess if ProtWave-VAE can perform semi-supervised learning for reshaping the latent space concerning fitness or function while retaining generative capacity, we compare an unsupervised ProtWave-VAE to a similar architecture with an additional regression decoder that samples from $z$ and predicts fitness $y$. The Chorismate mutase dataset is an ideal protein dataset as it includes enzyme sequences $x$ from a specific protein family and fitness measurements $y$ obtained by Russ et al. [13].

For the unsupervised learning architecture, we employed a model architecture similar to the one described above and optimized over comparable hyperparameters based on the negative log-likelihood on the hold-out set. In this instance, the training set consists of natural homologs, while the hold-out set contains synthetic designs. The final hyperparameter optimization values are: z_dim=4, C_out=256, num_fc=0, wave_hidden_state=64, head_hidden_state=512, num_dil_rates=8, NLL_weight=1, KL_weight= 0.95, and MMD_weight=10. The hyperparameter optimization search results can be found https://github.com/PraljakReps/ProtWaveVAE/tree/main/Pfam_analysis/outputs/hp_optimization/CM. The optimizer is Adam with a learning rate of $1 \times 10^{-4}$, 300 epochs, and a batch size of 512. The model was trained on unaligned sequences.

For semi-supervised learning, the gated dilated encoder and latent-conditioned WaveNet decoder architecture utilize the previously described hyperparameter optimized unsupervised architecture. We then introduce a discriminative decoder $p_\omega(y|z)$ with a depth of `disc_num_layers`=2, a linear layer width of `hidden_width`=10, and `p`=0.3. Here, we only implement a regression decoder and omit a classification decoder from `Decoder_re`. Next, we add a mean-squared error loss between the relative enrichment (r.e.) fitness prediction and ground truth to the unsupervised loss objective, applying a prefactor weight of 1. The results presented in Chorismate mutase (CM) inference and protein sequence generation for unsupervised and semi-supervised architectures use the above hyperparameters. Pretrained weights for these two models can be found https://github.com/PraljakReps/ProtWaveVAE/tree/main/Pfam_analysis/outputs/train_sess/pfam/CM.

#### 2.7 Model architecture and hyperparameter optimization for SH3 design task

To evaluate the ability of ProtWave-VAE to design functional sequences using semi-supervised learning and alignment-free sequence inference, we tasked the model with designing functional *in vivo* SH3 domains in *Saccharomyces cerevisiae*. The model consists of a gated dilated encoder $q_\phi(z|x)$, a latent-conditioned WaveNet decoder $p_\theta(x|z)$, and a discriminative decoder that includes a regression and classification path $p_\omega(y|z)$. The regression path samples $z$ and predicts continuous values $y_{re}$, which represent the relative enrichment (r.e.) scores that measure the functionality within the Sho1 osmosensing assay using next-generation sequencing. The classification path is independent of the regression path and samples $z$ to predict class probabilities $y_c$ using a final sigmoidal function layer. The two classes are functional (sequences with r.e. $\geq 0.5$) versus non-functional (sequences with r.e. $< 0.5$) sequences. The loss objective for classification is binary cross-entropy, while the loss objective for regression is mean-squared error. We performed hyperparameter optimization based on good negative log-likelihood, mean-squared error, and binary cross-entropy values, while maintaining excellent max-mean discrepancy regularization values. Based on the metrics and the procedure described in the previous sections for hyperparameter optimization, the final hyperparameter optimization configuration is as follows: (1) `z_dim`=6, (2) `C_out`=128, (3) `num_fc`=2, (4) `disc_num_layers`=2, (5) `hidden_width`=10, (6) `p`=0.4, (7) `whs`=256, (8) `hhs`=512, (9) `num_dil_rates`=12, (10) `NLL_weight`=1.0, (11) `KL_weight`=0.99, (12) `MMD_weight`=10.0, and (13) `gamma_weight`=1.0. The sequence data is alignment-free, and the optimizer implemented is Adam with a learning rate of $1 \times 10^{-4}$. We used a batch size of 1024 and trained for 200 epochs. The results of the hyperparameter optimization can be found at https://github.com/PraljakReps/ProtWaveVAE/tree/main/SH3_design_project/outputs/SH3_task/hp_optim.

We conducted stratified cross-validation instead of vanilla k-fold cross validation because the dataset contained a larger number of nonfunctional sequences than functional sequences. For stratified cross-validation, we trained five different train/validation configurations while monitoring the negative log-likelihood, mean-squared error loss, binary cross entropy loss, and max-mean discrepancy regularization loss, as well as classification performance metrics such as precision, recall, and F1 score on the validation set. The results, which can be found https://github.com/PraljakReps/ProtWaveVAE/tree/main/SH3_design_project/outputs/SH3_task/CV, show that the values are consistent and good over the five stratified train/validation splits. With the hyperparameter configuration and training configuration described above, ProtWave-VAE was trained on the entire dataset before generating and designing novel sequences for experimental testing. The pretrained weights can be found https://github.com/PraljakReps/ProtWaveVAE/tree/main/SH3_design_project/outputs/SH3_task/final_model

### 3 Supplementary Methods

#### 3.1 Random mutagenesis of SH3 sequences

The goal of this study was to enhance the functionality of a weak binding natural ortholog (subgroup III) and a weak binding natural hof1 paralog (subgroup IV) by using N-terminus plus latent conditioning and inpainting the C-terminus missing region. To avoid elevating functionality based on random chance, the generative model's performance was compared with that of random mutagenesis. To achieve this, we mutated the same C-terminus region of the weak ortholog and weak paralog randomly, while maintaining the same novelty as the designed inpainted sequences. The random mutagenesis sequences were substituted with amino acids belonging to the reference sequence at random sites along the C-terminus design region until we matched the minimum Levenshtein distance that corresponds to the generative designs. By doing so, we ensured that the random mutagenized sequences were mutated along the same inpainted region as the designs, while retaining the same diversity and novelty as the synthetic generative designs. The results showed that for weak binding ortholog subgroup IV and weak binding paralog group V, the random mutagenized sequences had matching edit distances to the design's minimum Levenshtein distance to the original weak binding natural paralog or ortholog, respectively. The matching edit distances were obtained when the N-terminus conditioning was 25%, 50%, or 75%. Figure S1A and B show the corresponding results for the weak binding ortholog and paralog, respectively.

### 4 Supplementary Results

#### 4.1 Pfam task: ColabFold structure prediction analysis of design sequences

A crucial evaluation of the ProtWave-VAE involves determining the extent to which the model can discern biologically relevant representations without receiving any annotations. We observed that the latent space sorts training sequences into phylogenetic groups and functional subclasses for S1A serine protease and beta-lactamase protein families (Figure S2A). The following assessment involves confirming if ProtWave-VAE can sample from this specific biologically significant latent space and produce new artificial sequences indistinguishable from the training set.

To evaluate the model's generative capabilities, we created alignment-free sequences, predicted associated tertiary structures utilizing ColabFold [8], and assessed if predictions mirrored the tertiary structures of natural homologs. For DHFR, S1A proteases, and lactamase protein families, we sampled 100 latent vectors $z$ from an isotropic Gaussian distribution and employed the ProtWave-VAE autoregressive decoder $p_\theta(x|z)$ to generate artificial sequences. Subsequently, we predicted the tertiary structure of each artificial sequence and calculated the TMscores and heavy-atom root mean squared distances (RMSDs). The anticipated tertiary structure with maximum, median, and minimum TMscores for the DHFR, S1A protease, and lactamase protein families are displayed in Figure S2B. We observed that the structure corresponding to the median TMscore accurately reflects the natural homolog's tertiary structure.

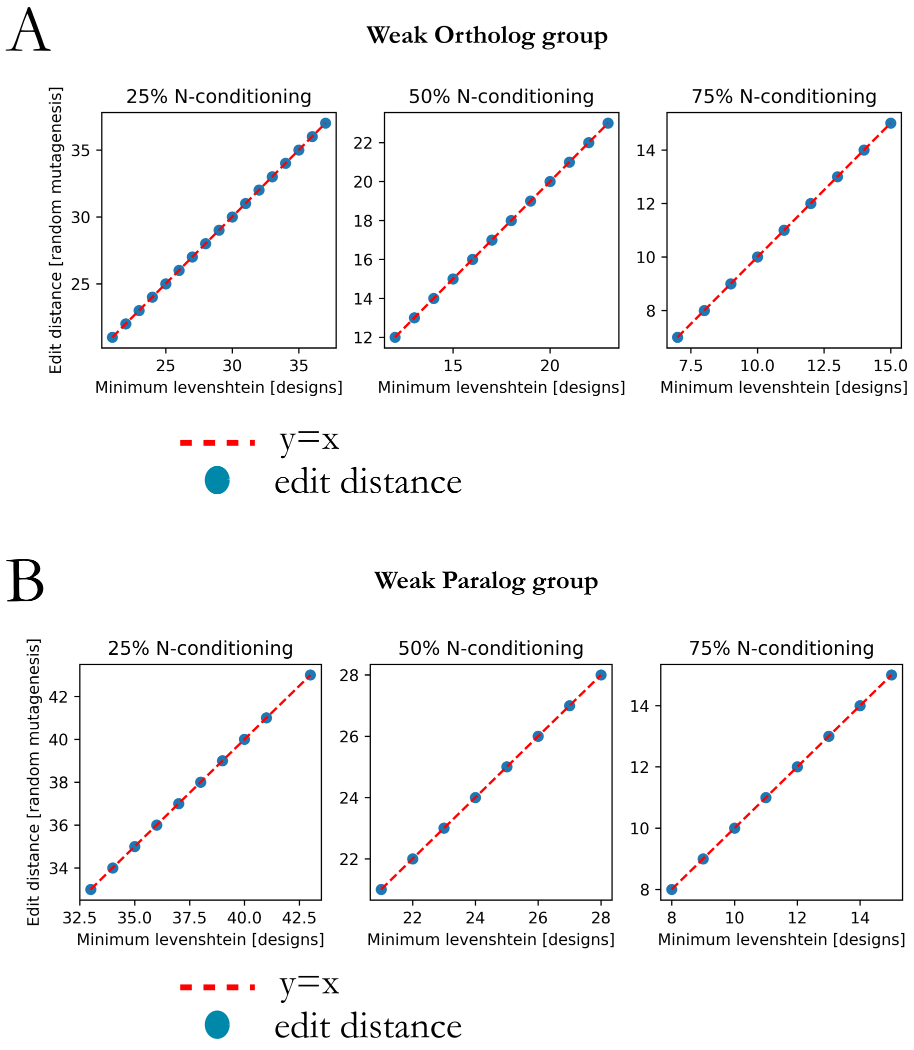

Figure S1: Random mutagenesis control for C-terminus diversification of protein designs with N- terminus plus latent conditioning. (A) The edit distance – number of random substitutions along the remaining C-terminus region of the reference weak binding ortholog – is shown on the y-axis for 25%, 50%, and 75% N-terminus conditioned subgroups. The x-axis shows the minimum Levenshtein distance between the designed sequences using N-terminus plus latent conditioning generative approach and weak binding natural ortholog sequence. (B) Similarly, the edit distance –number of random substitutions along the remaining C-terminus region of the reference weak binding paralog – is shown on the y-axis for 25%, 50%, and 75% N-terminus conditioned subgroups. The x-axis shows the minimum Levenshtein distance between the designed sequences using N-terminus plus latent conditioning generative approach and weak binding natural paralog sequence.

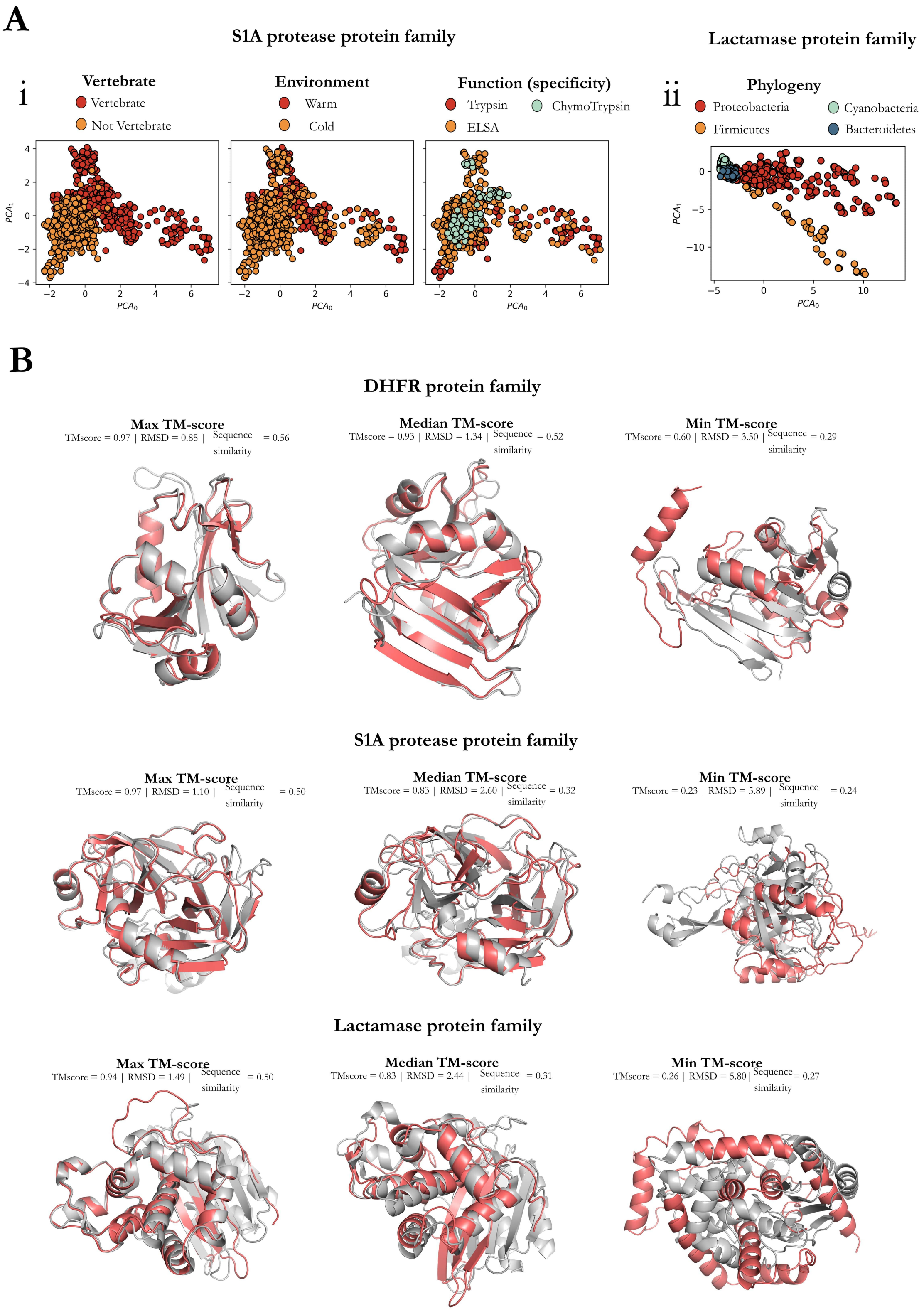
A
S1A protease protein family
Lactamase protein family
i
Vertebrate
Vertebrate
Not Vertebrate
Environment
Warm
Cold
Function (specificity)
Trypsin
ChymoTrypsin
ELSA
ii
Phylogeny
Proteobacteria
Cyanobacteria
Firmicutes
Bacteroidetes
$PCA_0$
$PCA_1$
B
DHFR protein family
Max TM-score
TMscore = 0.97 | RMSD = 0.85 | Sequence similarity = 0.56
Median TM-score
TMscore = 0.93 | RMSD = 1.34 | Sequence similarity = 0.52
Min TM-score
TMscore = 0.60 | RMSD = 3.50 | Sequence similarity = 0.29
S1A protease protein family
Max TM-score
TMscore = 0.97 | RMSD = 1.10 | Sequence similarity = 0.50
Median TM-score
TMscore = 0.83 | RMSD = 2.60 | Sequence similarity = 0.32
Min TM-score
TMscore = 0.23 | RMSD = 5.89 | Sequence similarity = 0.24
Lactamase protein family
Max TM-score
TMscore = 0.94 | RMSD = 1.49 | Sequence similarity = 0.50
Median TM-score
TMscore = 0.83 | RMSD = 2.44 | Sequence similarity = 0.31
Min TM-score
TMscore = 0.26 | RMSD = 5.80| Sequence similarity = 0.27

Figure S2: ProtWave-VAE infers meaningful biological representations on alignment-free protein families. (A) Principal component analysis (PCA) projections of the inferred latent spaces for the S1A proteases and beta-lactamase families are presented. The unsupervised model of the S1A protease family disentangles homologs within the inferred latent space based on vertebrate, environmental conditions, and functional specificity. For the beta-lactamase family, the model disentangles homologs in the inferred space in terms of the phylogeny. (B) To test the generative capacity of ProtWave-VAE, we randomly sampled 100 latent vectors z for each protein family from a normal distribution $\mathcal{N}$(0,I), corresponding to the latent prior. Using a computational structure prediction workflow (ColabFold + TMalign), we predicted each structure of the sample sequences and compared the predicted structure against a natural homolog that defines the corresponding protein family. We retrieved TMscores and root-mean-square distance (RMSD) scores. The structure predictions of ProtWave-VAE novel design sequences (red) for DHFR, S1A protease, and lactamase are visualized with the alignment of maximum, median, and minimum TM-score synthetic sequences against the natural reference homolog structure (grey).

#### 4.2 SH3 design task: experimental results

The relative enrichment (r.e.) score provides a quantitative measurement of the degree to which our designed SH3 domains are functional *in vivo* and capable of activating a homeostatic osmoprotective response. The assay shows good reproducibility in independent trials ($R^2$ = 0.94, $\rho_{pearson}$=0.97, n=1002, $p < 10^{-307}$, Figure S3).

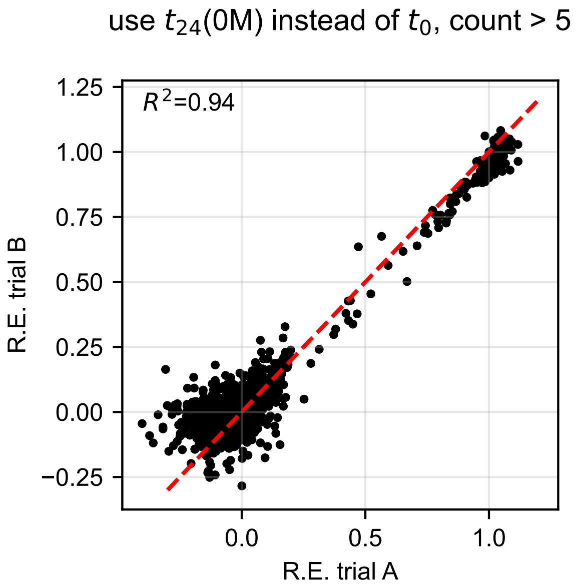

Figure S3: Validation of the high-throughput select-seq assay. A scatter plot of the enrichment score relative to wild-type *r.e.* for two independent ($n$ = 1002 including standard curve mutants) trials of the select-seq assay under the same experimental conditions. The position of the wild-type Sho1 sequence is normalized to be at $(1, 1)$ and of the null allele is at $(0, 0)$. The red dashed line indicates the identity trace. The data shows that the select-seq assay shows good reproducibility between independent runs ($R^2 = 0.94$; $\rho_{Pearson} = 0.97, n = 1002, p < 10^{-307}$)

[3] Junyoung Chung, Caglar Gulcehre, KyungHyun Cho, and Yoshua Bengio. Empirical evaluation of gated recurrent neural networks on sequence modeling. *arXiv preprint arXiv:1412.3555*, 2014.

[16] Aaron van den Oord, Sander Dieleman, Heiga Zen, Karen Simonyan, Oriol Vinyals, Alex Graves, Nal Kalchbrenner, Andrew Senior, and Koray Kavukcuoglu. Wavenet: A generative model for raw audio. *arXiv preprint arXiv:1609.03499*, 2016.

[17] Aaron Van den Oord, Nal Kalchbrenner, Lasse Espeholt, Oriol Vinyals, Alex Graves, et al. Conditional image generation with pixelcnn decoders. In D. Lee, M. Sugiyama, U. Luxburg, I. Guyon, and R. Garnett, editors, *Advances in Neural Information Processing Systems 29 (NIPS 2016)*, pages 4790–4798, 2016.

[18] Nicholas C Wu, Lei Dai, C Anders Olson, James O Lloyd-Smith, and Ren Sun. Adaptation in protein fitness landscapes is facilitated by indirect paths. *Elife*, 5:e16965, 2016.

[19] Ran Zhang, Lin Cao, Mengtian Cui, Zixian Sun, Mingxu Hu, Rouxuan Zhang, William Stuart, Xiaochu Zhao, Zirui Yang, Xueming Li, Yuna Sun, Shentao Li, Wei Ding, Zhiyong Lou, and Zihe Rao. Adeno-associated virus 2 bound to its cellular receptor AAVR. *Nature Microbiology*, 4(4):675–682, 2019.
